# Modulation of signal response by small RNAs through feed-forward loops

**DOI:** 10.64898/2025.12.22.696124

**Authors:** Yang-Chi-Dung Lin, Yigang Chen, Ting-Syuan Lin, Shenghan Huang, Xiao-Xuan Cai, Jie Ni, Liping Li, His-Yuan Huang, Hsien-Da Huang

## Abstract

In response to environmental changes, bacteria have evolved sophisticated regulatory networks that incorporate small RNAs (sRNAs) and transcription factors (TFs) to fine-tune cellular physiology. Both sRNAs and TFs modulate gene expression, but the former function via post-transcriptional mechanisms, while the latter act at the transcriptional level. However, it remains unclear why both regulatory layers are conserved through evolution, rather than one being sufficient. Here, we experimentally identified that CRP, a global regulator, regulates 25 small RNAs (sRNAs) in *Escherichia coli*. Interestingly, CRP also controls 80% of the target genes of these sRNAs. This architecture led us to identify 34 novel sRNA-mediated feed-forward loops (sFFLs)—circuits where CRP regulates both an sRNA and its target—within the CRP regulon. Quantitative PCR analysis of 16 such sFFLs revealed that each type possesses a distinct cAMP dose-response profile, suggesting that different sFFL structures embody unique regulatory logic. Specifically, coherent and incoherent type 3 and 4 sFFLs exhibit broader dynamic ranges in their dose-response compared to open-loop controls. Coherent and incoherent type 1 and 2 sFFLs appear more energy-efficient. Altogether, we propose that sRNAs cooperate with TFs through sFFLs to optimize both the energy efficiency and the diversity of signal response profiles. Therefore, sRNAs serve as critical components integrating transcriptional and post-transcriptional networks across diverse cellular pathways.

## INTRODUCTION

Living in an ever-changing environment, organisms must detect changes in different parameters to coordinate their network system. Therefore, organisms have evolved sophisticated mechanisms that can simultaneously transduce and calculate. When the existence of a specific circuit in a network is more frequently than in a random network, this circuit is called a network motif (1). Key components of these motifs involve the interplay between transcriptional and non-transcriptional regulation, utilizing mechanisms such as kinase-mediated phosphorylation (2), protease-mediated proteolysis (3), and RNA-mediated regulation (4). In bacteria, RNA-mediated regulation predominantly involves small RNAs (sRNAs). sRNAs can post-transcription modulate target genes by either enhancing or repressing transcription and translation (5). Most sRNAs base-pairdirectly with their target mRNAs, especially in the five prime untranslated regions (5’-UTR), to block the ribosome binding site, inhibit translation, and accelerate the degradation of targets, or to remodel secondary structure in the 5’-UTR to facilitate translation (5–7). Some sRNAs, however, work through binding proteins. These protein-binding sRNAs act to antagonize the regulatory function of their cognate proteins by competing with substrate RNA molecules to bind these proteins. For example, in *E. coli*, the sRNAs CsrB and CsrC repress CsrA protein activity by presenting multiple binding sites that compete with other RNA targets (8,9). Similarly, the sRNA GlmY is reported to stabilize the transcript of another sRNA, GlmZ, by sequestering the processing protein YhbJ (10).

Nevertheless, post-transcriptional regulation cannot do without transcriptional regulation to exert all kinds of cellular events. Like sRNAs, many proteins that act as transcription factors can also regulate the abundance of their target genes, but through mechanisms distinct form sRNA. They act trough interaction with RNA polymerase, and other transcriptional factors to initiate or hinder the transcription of target genes (11).

Although sRNAs have been reported to participate in diverse cellular activities such as mobility (12), biofilm formation (12) and metabolic regulation (13,14), the biological significance of sRNA-mediated regulation remains under debate. The advantage of using sRNA for energy conservation seems unlikely, as using TFs to regulate gene expression saves more energy compared to using sRNAs (15). In general, TFs are relatively more stable than sRNAs, and they do not degrade when binding promoters of target genes (16). On the other hand, sRNA works through a non-catalytic process, where sRNA and its target are both degraded after base pairing (17). Therefore, the number of TF molecules required for repressing target gene expression is significantly smaller than that of sRNA (15). This raises a long-standing evolutionary question: What advantages might cells gain if they continuously use sRNAs instead of proteins?

The answer remains under debate. Some evolutionary studies regard sRNAs as relics of the early RNA world, preserved through mechanisms requiring specific nucleic acid recognition (18–20). Other studies focus on the kinetic differences between sRNAs and proteins, claiming that sRNAs are more suitable for switching cellular states in stress responses (16,21–23). Given that sRNAs can base-pair with their targets immediately after transcription, their regulatory effect is faster than that of TFs, which still require the translation process. However, this explanation is undermined by the facts that the expressions of sRNAs is itself controlled by TFs and, that environmental stimuli (such as pH, temperature, and osmolality) and the availability of small molecules (such as cAMP, oxygen, iron, and lactose) are usually directly sensed and responded to by TFs or by their cognate sensors (24–30).

To gain deeper insight into this regulatory interplay, we employed a systemic approach to identify cAMP receptor protein (CRP)-regulated genes in E. coli. Unexpectedly, we found that 33% (25 out of 80) of sRNAs are regulated by CRP, and 85% (41 out of 48) of the targets of these sRNAs are also regulated by CRP. Thus, the CRP regulon is enriched with recurrent circuits where CRP regulates both an sRNA and its target gene.

These circuits form feed-forward loops (FFLs), a dominant motif in three-gene networks. In a generic FFL, a primary regulator (here, CRP) controls an output target gene both directly and indirectly. When the indirect path is mediated by an sRNA, these are called “sRNA-mediated feed-forward loops (sFFLs)” (21), analogous to microRNA-mediated feed-forward loops (FFLs) that pervade the regulatory networks in mammalian cells (31,32). In the past decade, the function of transcriptional FFLs (encompassing only transcriptional regulation) has been studied by computational simulation and in vivo experiments (33,34). According to U. Alon’s definition, these motifs are categorized into coherent and incoherent FFLs, depending on whether the regulatory effect of two arms is synergistic or antagonistic (33). Each type of FFL can play a different role, including noise buffering (35), fold-change detection (36), generating periodic impulses (37), accelerating response time (34), as well as a filter-like function, limiting the system’s response to input within a narrow range (37). However, comprehensive research on sFFLs remains scarce (38), and theoretical models are lacking.

In this study, we utilized these CRP-regulated circuits to investigate the signal response of sFFLs. The cAMP dose-responses of 16 different sFFLs were quantified by real-time PCR (39). Interestingly, each type of sFFLs exhibited distinct dose-response profiles, implying that the specific topology and interaction parameters deetermine the circuit’s behavior.

We proposed a model to estimate the dose-response of various FFLs, suggesting that the regulatory effect of sRNA is limites by the amount of both sRNA and target transcript; thus, each FFL has an optimized working dose range where the regulatory effect of sRNA reaches its peak. Altogether, we conclude that most of the sRNAs may work in concert with a transcription factor to modulate their targets through FFL motif, and that each FFL possesses a specific dose-response and working dose range. sRNA-mediated FFLs demonstrates the cross-talk between transcriptional and post-transcriptional regulation. Our study on sRNA-mediated FFLs may serve as a fundamental building block for understanding how transcriptional and post-transcriptional networks coordinate to determine complex cellular responses.

## RESULTS

### CRP-regulated sRNAs involve Feed-forward loops

To understand the biological role of sRNA, we first need to clarify whether most of the sRNAs work cooperatively with transcription factors or are independently regulated to distinct functional gene groups. Nevertheless, the transcriptional regulations of most sRNAs remain unclear. We focused on identifying CRP-regulated sRNA because CRP is estimated to regulate approximately one-tenth of the genes in E. coli, and CRP has been reported to regulate some sRNA, such as spf and *cyaR*, in *E. coli* (*40*). We developed a systematic approach (**Figure 1A**), which combines computational prediction and differential expression data to clarify how many sRNA and their targets are directly regulated by CRP at the transcriptional level (41–43). First, CRP binding sites on the promoter of all known sRNAs in *E. coli* were identified by an HMM program and then confirmed by EMSA (**Figure 2, data not shown**). Independently, a customized sRNA-focused array (GEO accession: GPL10535) was used to quantify the differential expression of sRNAs in *crp* deletion mutant and wild-type strains (**Table 1A and Table 1B**). Combining the evidence of the CRP binding site and differential expression, 25 sRNAs have been considered to be regulated by CRP directly (**Figure 1B**). In other words, about one-third of the sRNAs in *E. coli* are regulated by CRP, which is ten times the size of previously reported cases.

**Figure 1A.**
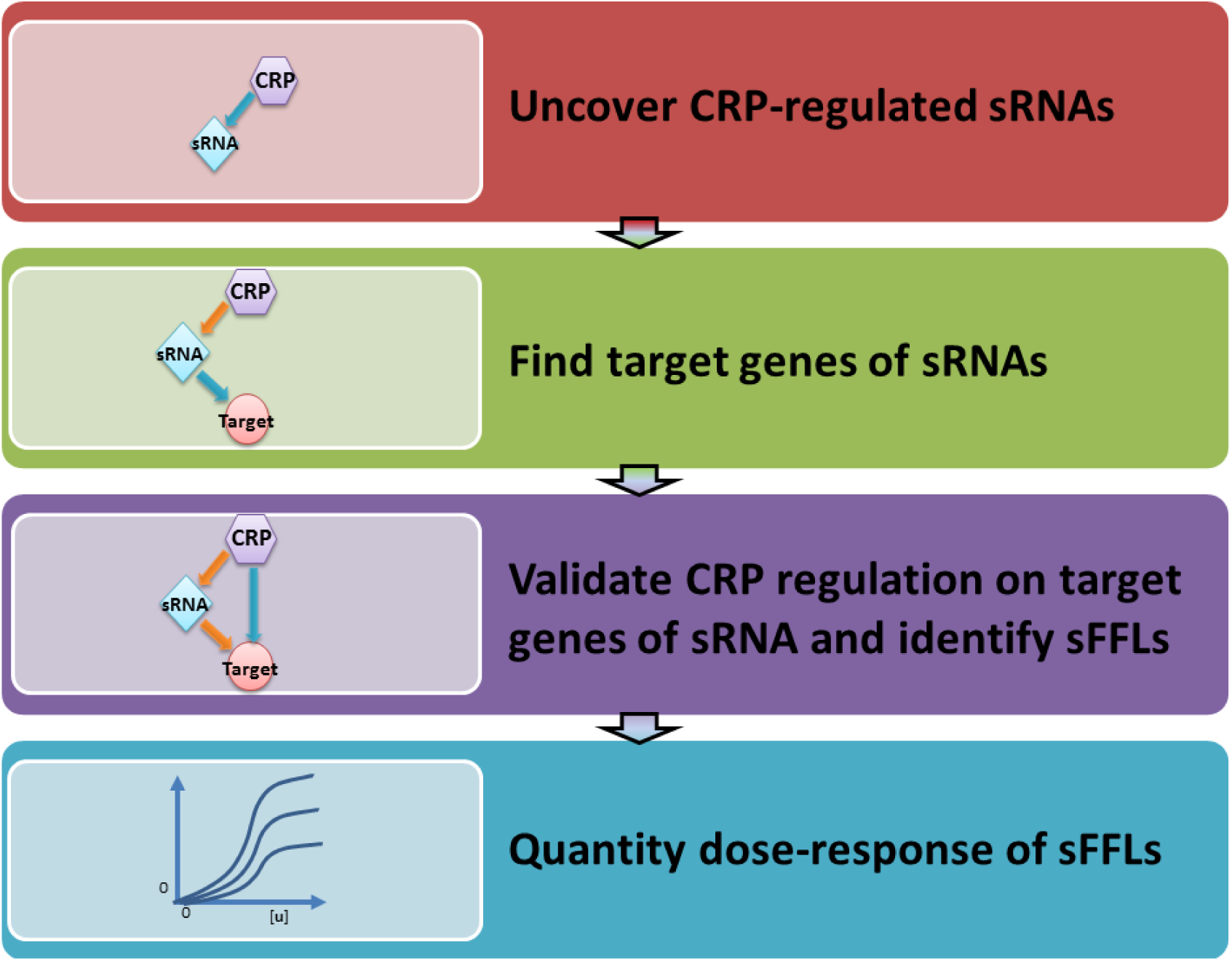
Steps of identifying signal response by modeling FFLs with CRP regulated sRNAs. First, CRP-regulated sRNAs were identified by predicting CRP binding sites in the sRNA promoters and measuring the differential expression of sRNAs between wild-type and *crp* deletion strains. Next, the regulatory effects of a CRP-regulated sRNA to its target genes were derived from literature review. If a target gene of CRP-regulated sRNA also has CRP binding site in promoter region and shows differential expression, a feed-forward loop comprising CRP, sRNA and a target gene is validated. The cAMP dose-responses of a sRNA was measured in wild-type strain, whereas that of its target genes were measured in wild-type and sRNA deletion strains. Finally, the relative expression profiles of the sRNA and its target genes were used as input to develop a mathematical model.

**Figure 1B.**
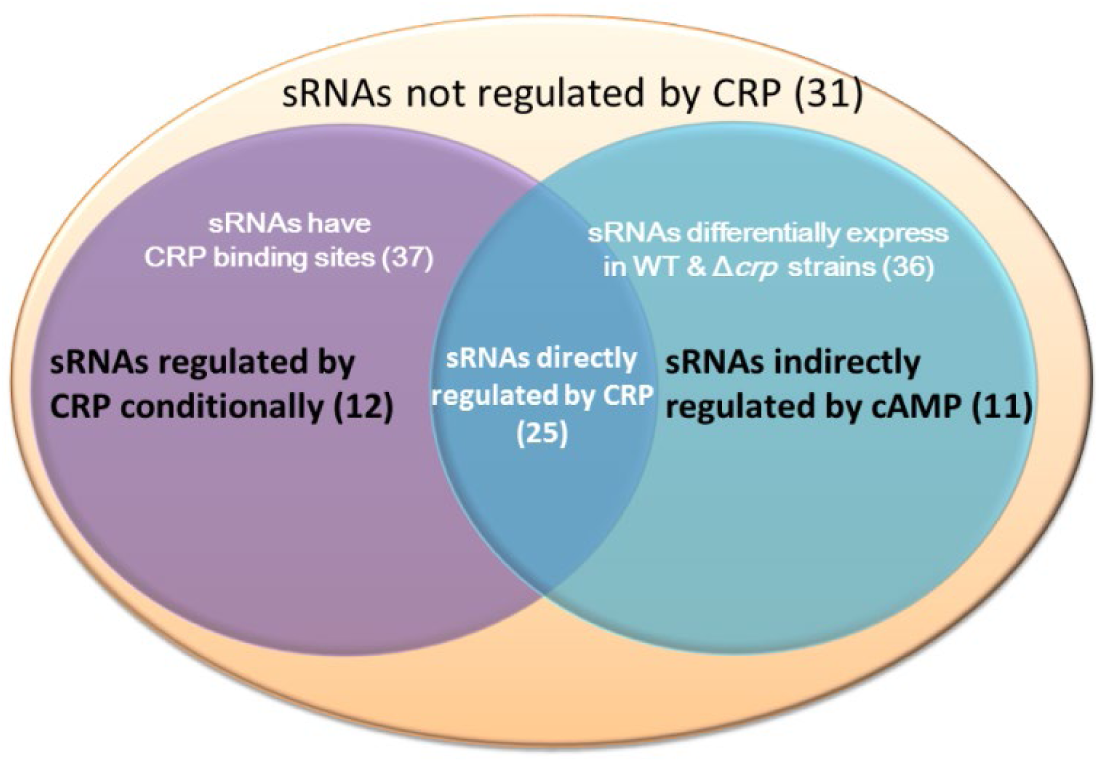
Regulatory effects of CRP on sRNAs in *E. coli*. In the two-circle Venn diagram, the purple circle encompass the sRNAs that have CRP binding sites but do not exhibit cAMP dose-response in our test condition, suggesting that CRP may dependents on other cofactors that express under specific conditions to regulate these genes. The blue circle includes all of the sRNAs that show detectable cAMP dose-response using microarray, as well as some sRNAs that show cAMP dose-response in specific growth conditions. The overlap area in this diagram denotes the sRNAs that have CRP binding sites and exhibit significant differential expression in wild-type and *crp* deletion strains.

**Figure 1C.**
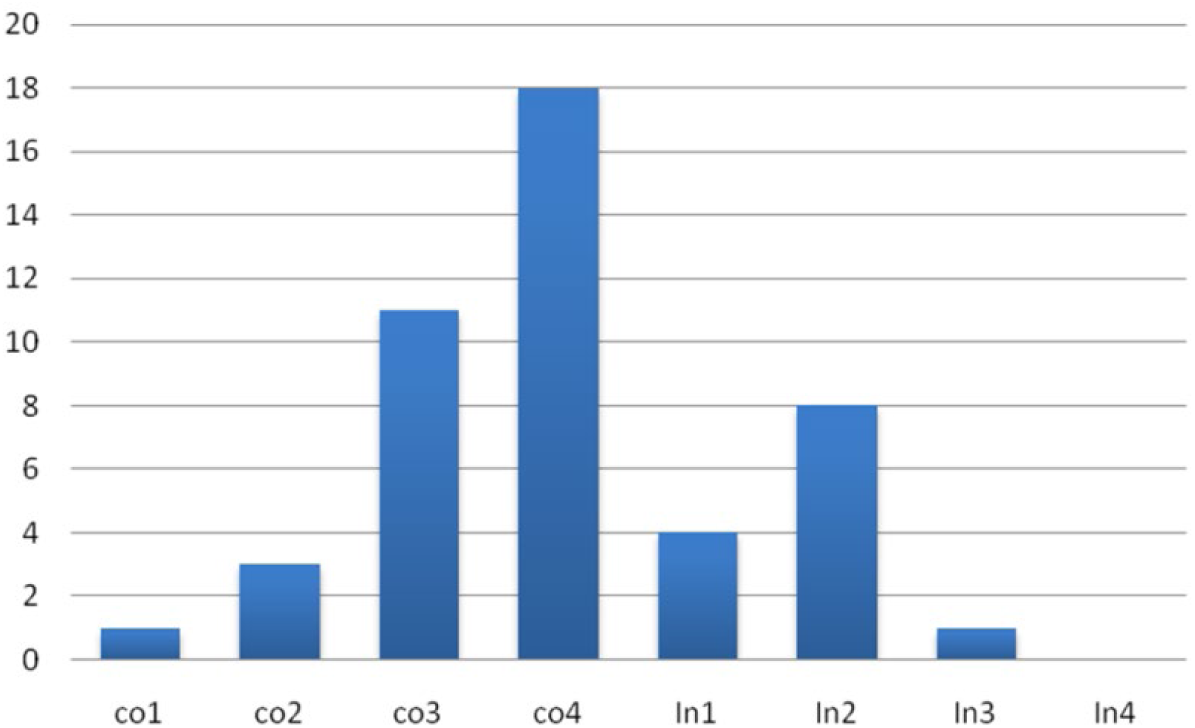
Frequency analysis of each type of sFFL in CRP regulon. All of the eight kinds of sFFL, except incoherent type 4 sFFL, were found in CRP regulon. The bar charts summarize the frequencies of each kind of sFFL that is regulated by CRP and a CRP-regulated sRNA in *E. coli*. The Y-axial represent the number of the sFFLs belong to a specific type.

**Figure 1D.**
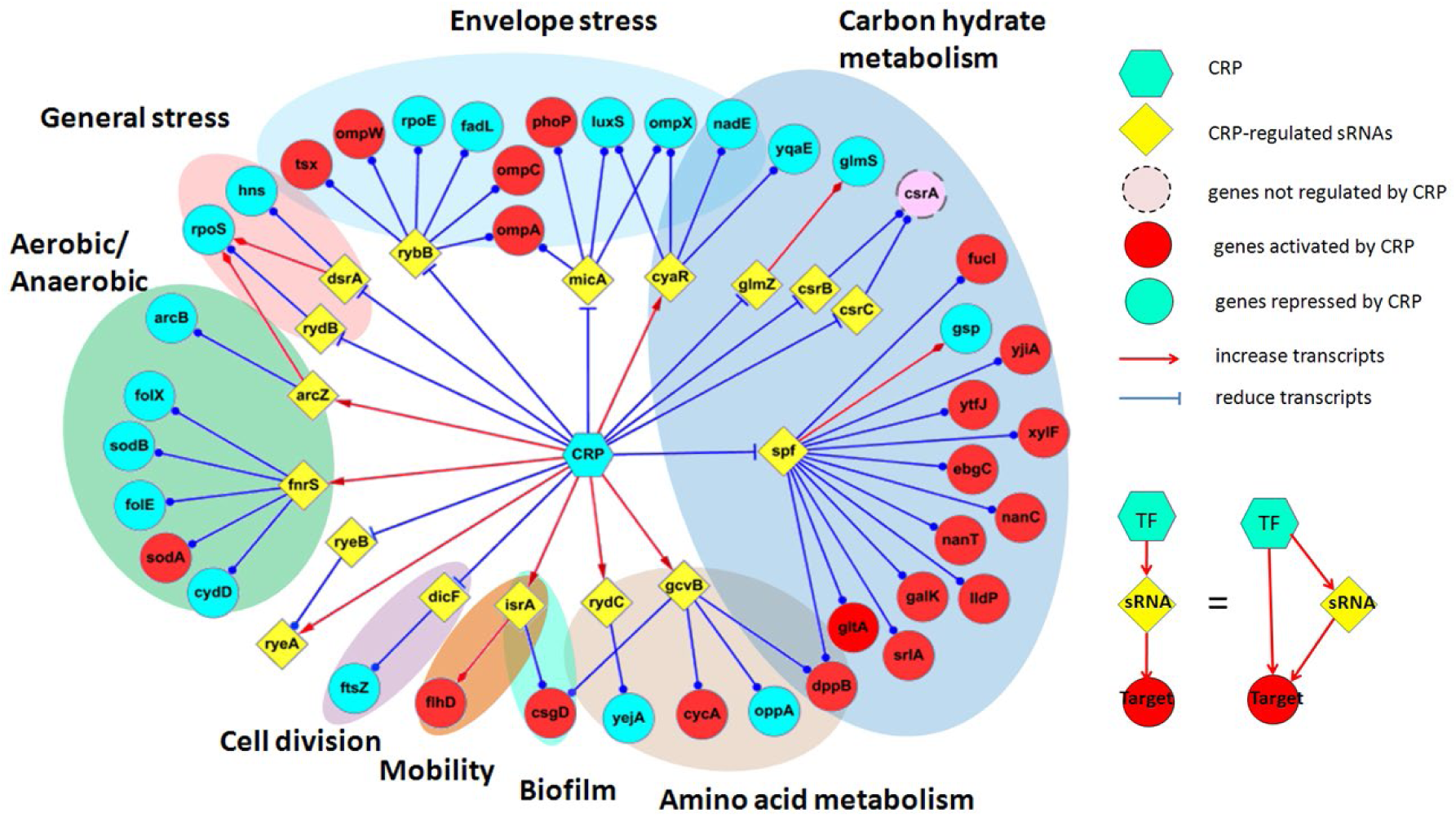
Functional network of CRP–regulated sFFLs. This diagram only shows the genes involve sRNAs-mediated FFL. The node shape represents three gene classes: cAMP receptor protein (octagon). sRNAs (rhombus), target genes of sRNAs and CRP (circle).Regulatory effects that enhance the abundance of target gene transcript are indicated by arrows (→), while those destabilize target gene transcript are illustrated as blunt ends (⊥). To simplify the picture, CRP-mediated regulations on sRNA target genes are indicated by the color of the circles. Blue circles indicate the gene is activated by CRP; whereas the red circles represent CRP repressed genes. The transcriptional regulations between CRP and sRNA target genes were measured under aerobic batch culture at 37C. The interaction information between sRNA genes and their targets were taken from sRNAMAp[reference], EcoCyc (Keseler et al., 2011), RegulonDB (Gama-Castro et al., 2011), NPInter (Wu et al., 2006)and sRNATarBase (Cao et al., 2010), detail reference about each interaction is available in Supplement table 1.

**Figure 2.**
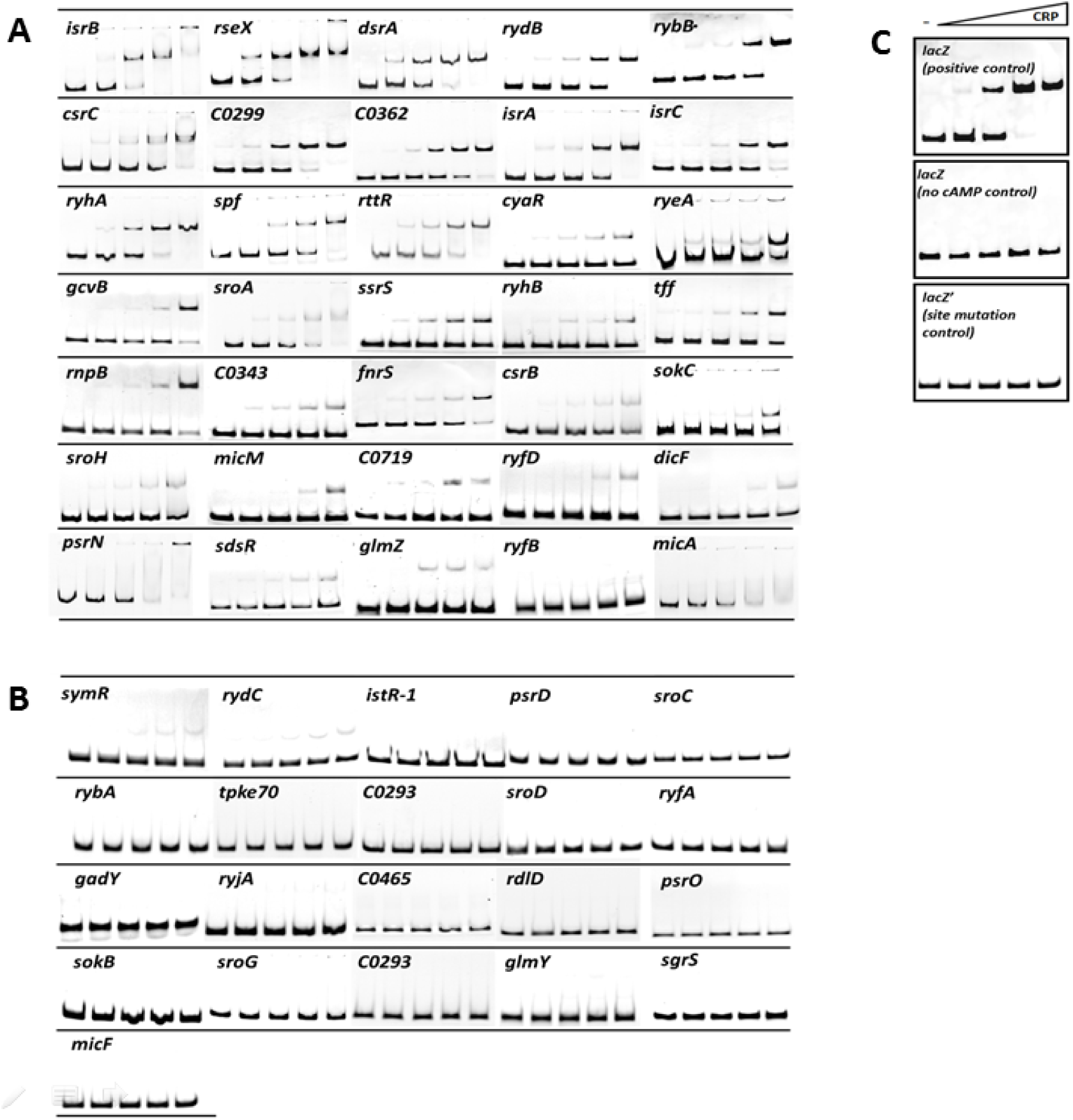
The cAMP-CRP binds to the promoter regions of the sRNAs. 56 computational predicted CRP binging sites scored above 4.9 were further validated by EMSA. The results show that the false negative rate is 3%. (A)The upper panel presents the EMSA results of predicted CRP binding site scored above 6.0, (B) whereas the lower panel shows binding site score between 4.9 to 6.0 with CRP binding sites. In general, the higher the prediction scores it, the lower CRP concentration needed to induce band shift. (C) Wild-type and CRP-binding site mutant *lacZ* promoter sequences served as positive and negative control, respectively.

**Table 1A.**
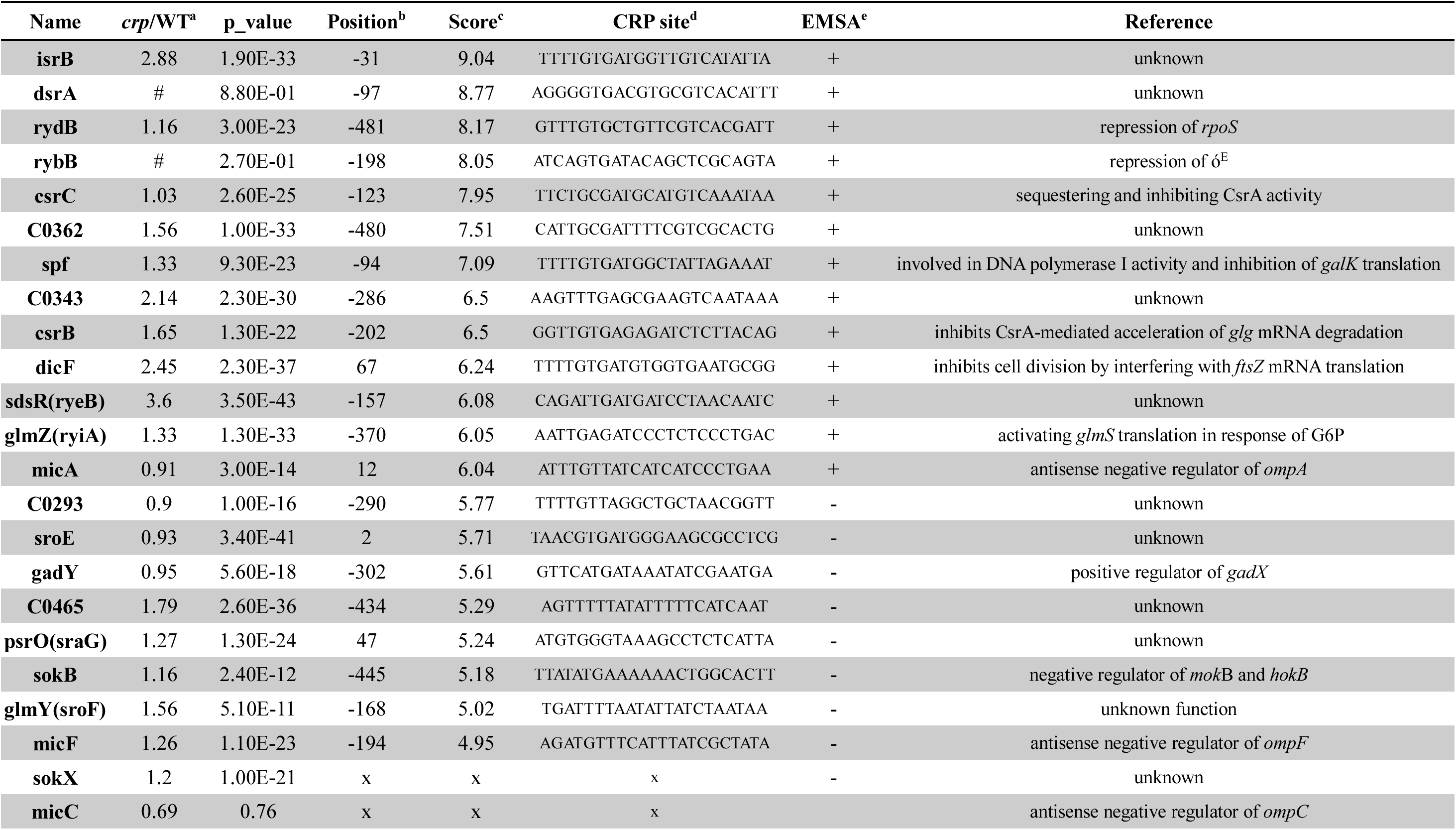
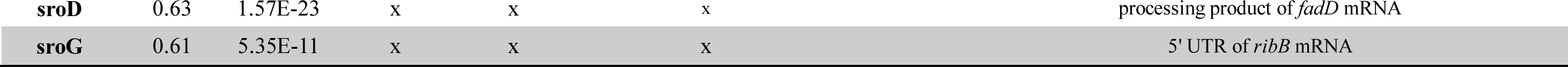
Collection of sRNAs up-regulated by CRP mutant in *Escherichia coli*.

**Table 1B.**
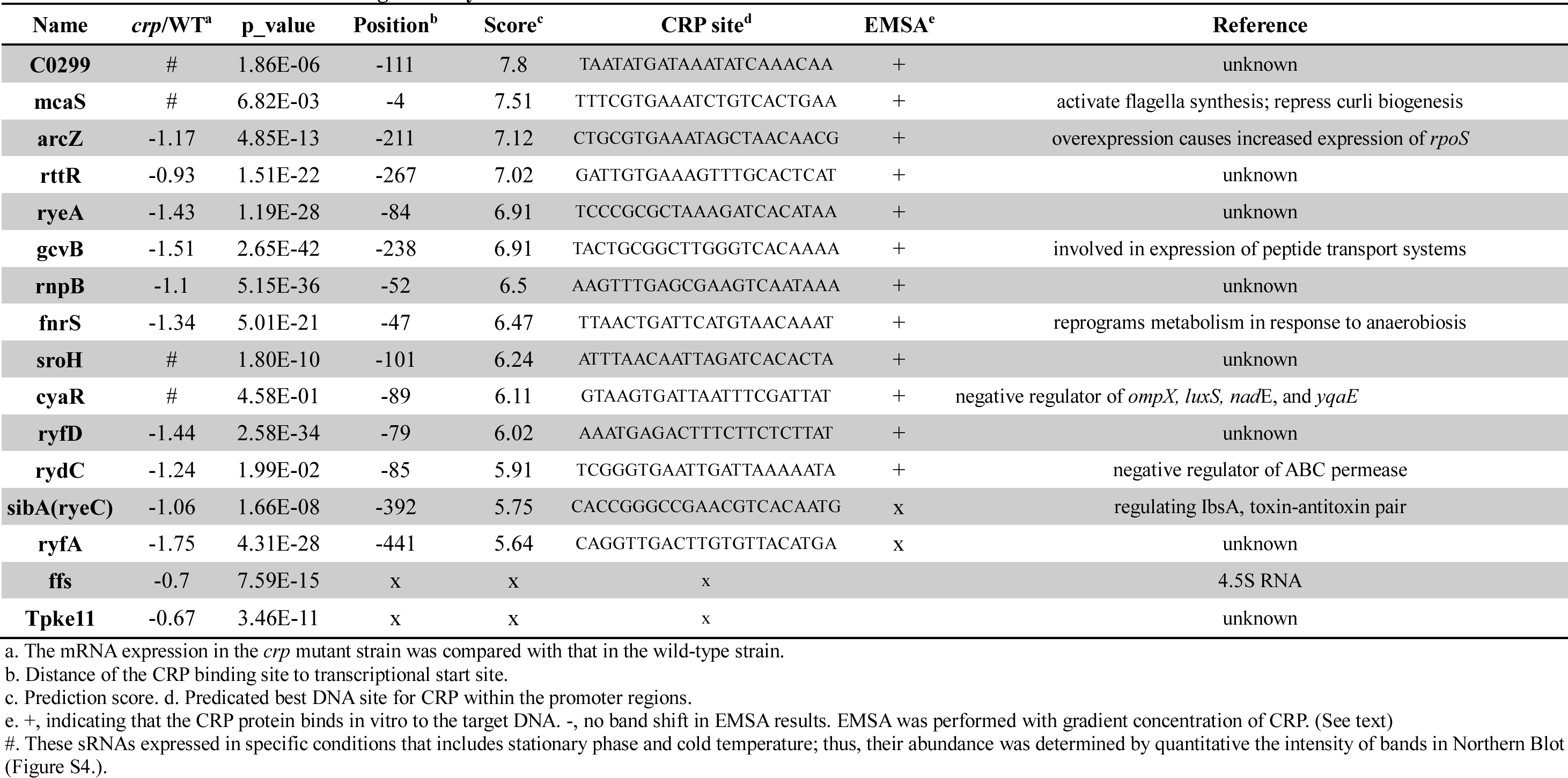
Collection of sRNAs down-regulated by CRP mutant in *Escherichia coli*.

Next, we investigated whether the target of those CRP-regulated sRNAs is also involved in the CRP regulon. Since sRNAs are non-coding, their function is dependent on their target genes. The interaction information between sRNA genes and their targets is experimentally determined target-binding region is taken from sRNAMAp (44), which is collected from EcoCyc (40), RegulonDB (45), NPInter (46) and sRNATarBase (47). Then, CRP binding sites on the promoter regions of sRNA target genes were predicted using the HMM program, and their differential expression in wild-type and *crp* deletion strains was validated by Real-time PCR (**Table 2**) and EMSA (**data not shown**). In addition to 39 sRNA target genes reported to be regulated by CRP, 22 novel target genes of CRP-regulated sRNAs were identified by our approach (**Supplementary File S1**). The frequency of an sRNA target with a CRP site is approximated 63% (51/81). Strikingly, the frequency is highest in sRNAs targets that regulated by CRP directly (83%, 40/48; hyper geometric test P <3*10^-47^) compared with that of the whole genome (4%) as well as that of sRNAs not regulated by CRP (46%, 7/15; hyper geometric test P<10^-7^) (40,45) (**Supplementary Table S1**). The dramatic contrast between these two groups indicated that CRP-regulated sRNA may cooperate with CRP to regulate their target genes. Thus, the regulation of their target genes is dependent on a type of FFL that comprises both transcriptional and posttranscriptional regulation, as such recurrent circuits are similar to microRNA-mediated Feed-forward loops, which are found in mammalian cells (31). These named them sRNA-mediated Feed-forward loops (sFFLs).

**Table 2.**
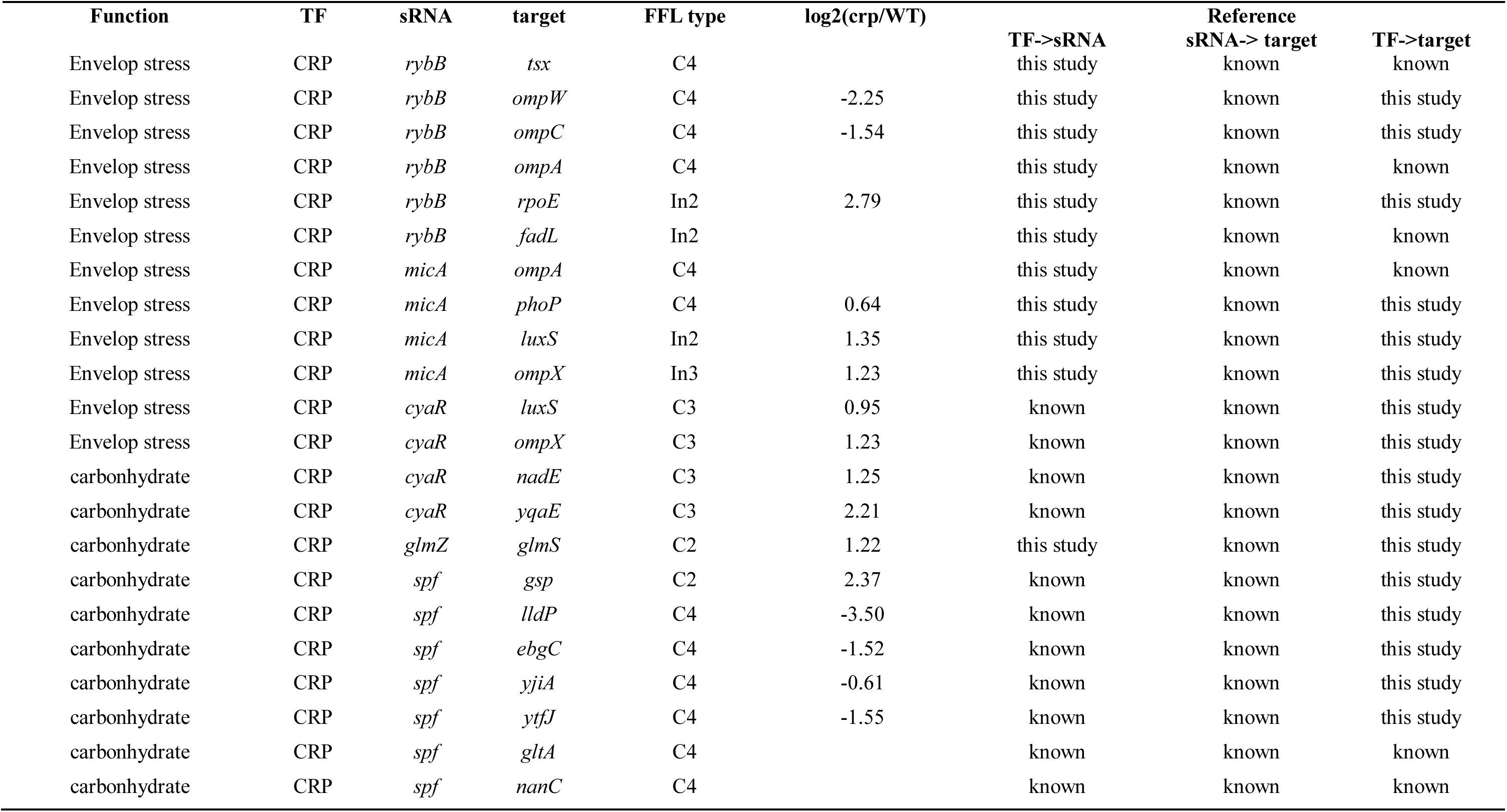
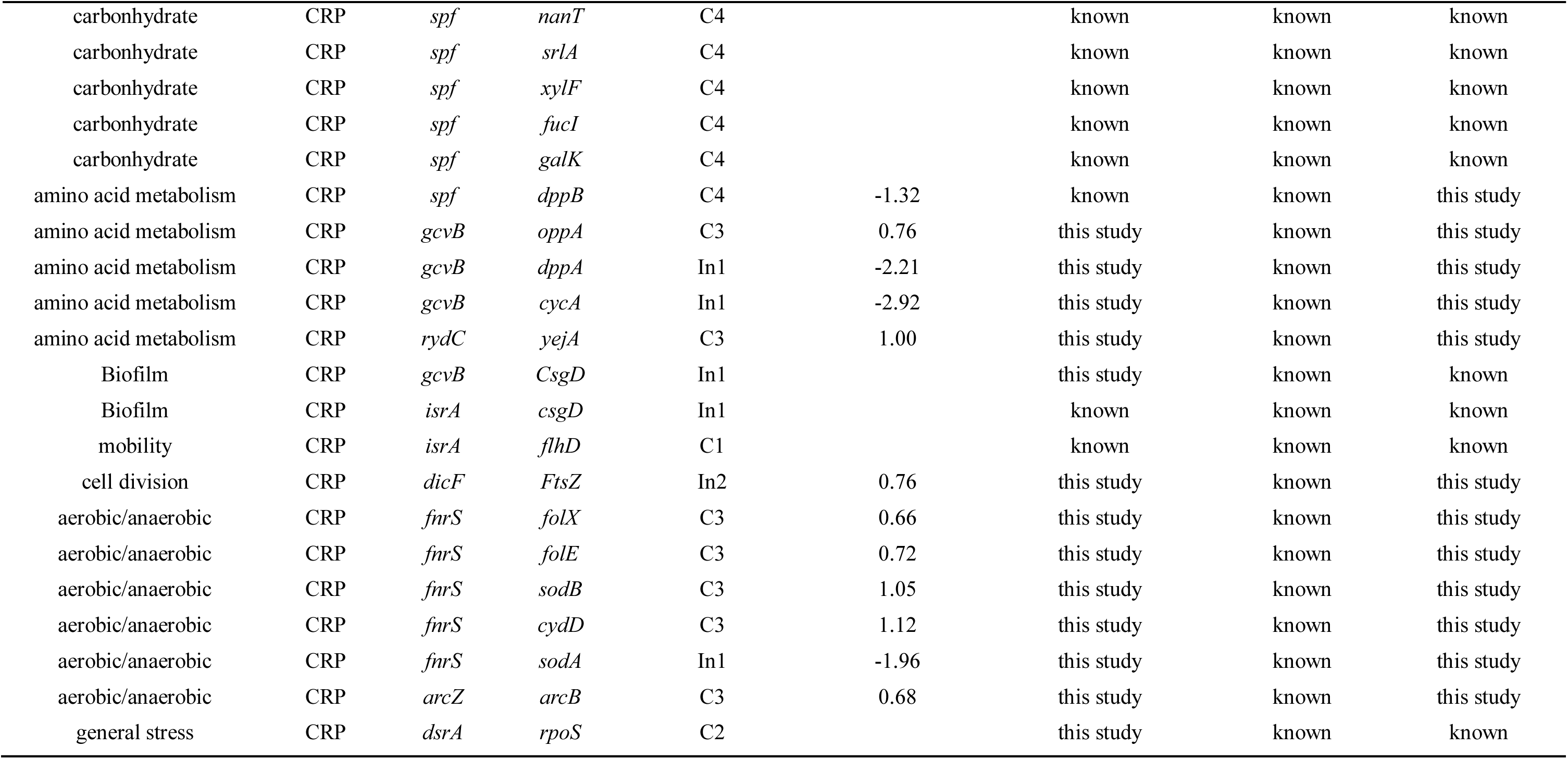
Collection of CRP regulated sRNA mediated FFL in *Escherichia coli*.

Summarizing the expression condition of sRNAs and their targets reveals that most of the CRP-regulated sRNA expressed during exponential phase: aerobic (ArcZ) (48) and anaerobic growth (FnrS) (49); biosynthesis of purine, pyrimidine, and amino acids (RydC and GcvB) (50,51); envelope stress response (RybB, MicA and CyaR) (52–54); carbohydrate metabolism (CsrB, CsrC, Spf, and CyaR) (9,53,55); general stress response (RydB and DsrA); cell division (DicF) (44); mobility (McaS); biofilm formation(McaS) (12). When all of these novel transcriptional regulation data were integrated with literature records, we found 35 novel feed-forward loops (**Supplementary File S2**). Except for incoherent type 4 FFLs, all kinds of s FFLs are found in the CRP-regulon (**Figure 1C**). As mentioned above, our new network reveals that CRP collaborates with numerous sRNAs to regulate a significant number of genes (**Figure 1D**). In this network, coherent type 4 is the dominant sFFL form, which differs from the previously reported distribution of FFLs with transcription regulation alone (37).

### sRNA-mediated feed-forward loops exhibit distinct dose-response profiles

To characterize the signal response properties of these motifs, we selected 16 sRNA-mediated FFLs covering seven out of the eight defined FFLs to conduct a dose-response assay (**Figure 3 and Supplementary Figure S1**). Since the DNA-binding affinity of CRP is directly dependent on intracellular cAMP levels, the expression of CRP targets functions as a response to cAMP concentration (56–61). To explicitly control activity of CRP, we utilized the *cyaA*, *tolC* double mutant strain, which is unable to synthesize or export cAMP. This allowed us to precisely modulate intracellular cAMP levels by exogenous addition and measure the relative expression of target genes in the presence or absence of specific sRNAs (61).

**Figure 3.**
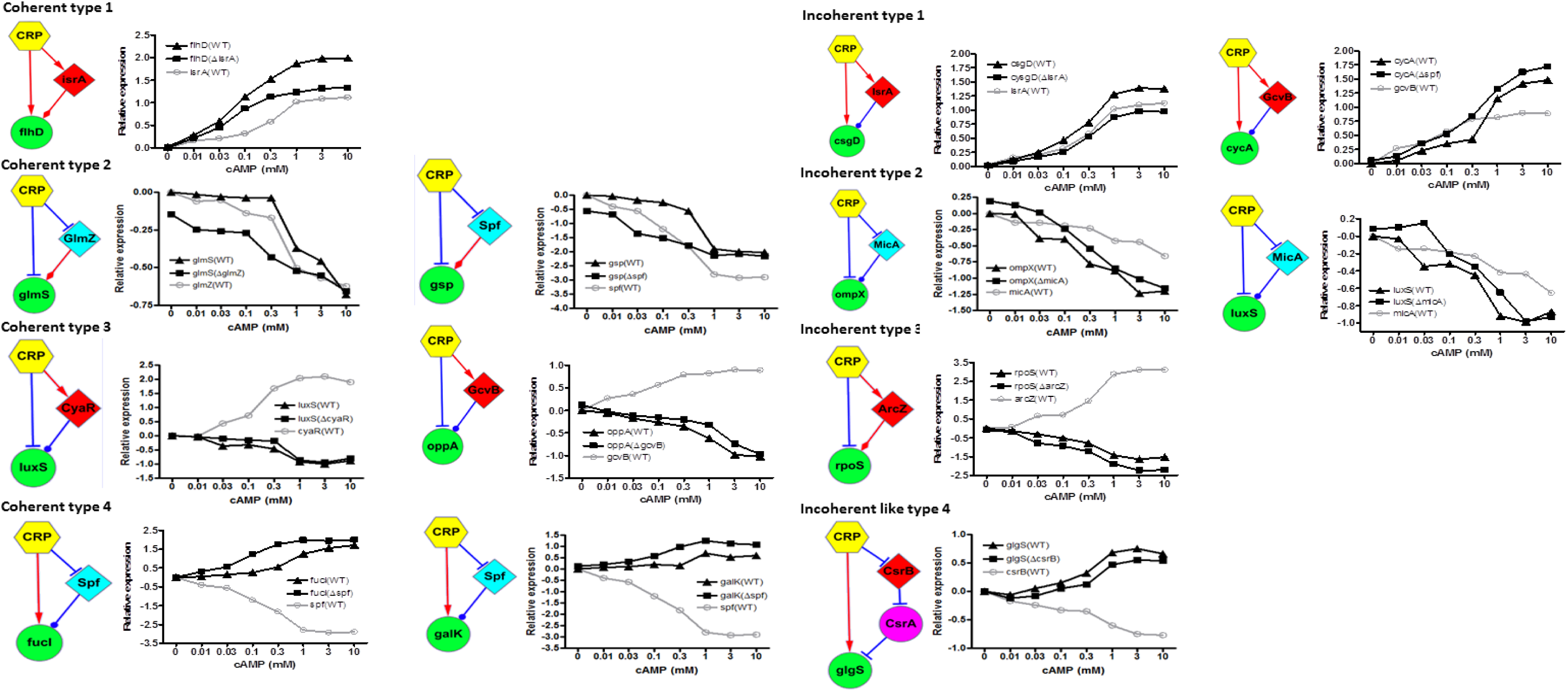
The cAMP dose-response curves of coherent and incoherent sFFLs. The expressions of sRNAs are draw in gray lines, whereas those of sRNA target genes are illustrated in black lines. The abundance of the *sRNA* (○) as well as its target genes in wild-type (▴) and sRNA mutant strains (▪) was determined by real-time PCR. The CsrAB system is not an incoherent type 4 FFL, because *csrB* act through repressed the function of *csrA* rather than direct base-pairing with its target. However, we treated it as an incoherent type 4-like FFL, because *csrB* cooperate with CRP to enhance the abundance of target gene. The *E. coli* BW25113 derivate Δ*cyaA* Δ*tolC* strain was referred to as the “wild-type” strain (WT) in this experiment, and all of the mutant strains were derivatives of this strain. The cells were grown in LB broth until OD600 reached 0.4. The cultures were treated with cAMP in various concentration for 10 minutes and then mixed with RNA stop solution and harvested to RNA extraction. Each point represents the average value of three independent experiments (P value <0.05). See Figure S1 for more dose-response data for different sFFLs.

We investigated whether the same kinds of FFLs select different profiles by measuring the responses of various FFL types, including two coherent Type 2 FFLs, two incoherent Type 1 FFLs, three coherent Type 3 FFLs, and four coherent Type 4 FFLs. Interestingly, each type of FFL appears to prefer a distinct dose-response profile. For example, either *glmZ* and *glmS* or *spf* and *gsp* can form a coherent type 2 FFL with CRP (**Figure 3**). Both pairs exhibited similar dose-response profiles, characterized by significant sRNA-mediated stability even at high cAMP concentrations. Conversely, the incoherent Type 1 FFLs formed by *isrA*/*csgD* or *gcvB*/*cycA* demonstrated sRNA-mediated repression that intensified with increasing cAMP concentrations (**Figure 3**). Altogether, these results suggest that the dose-response of each type of sFFL may be related to its topology, transcriptional regulation, and post-transcriptional regulation.

### CRP cooperates with the CsrABC system in an incoherent FFL

Although the feed-forward loop is defined as a three-gene pattern, we found that a sub-network encompassing four genes also behaves like an incoherent type 4 system. This pattern is formed by CRP, CsrA, CsrB, and *glgS* (**Figure 3**). CsrB and CsrC are protein-binding sRNAs (9,55). They contain multiple CsrA binding sites (18 in CsrB and nine in CsrC) and function to sequester the CsrA, preventing it from binding to other mRNAs (62). In most cases, the binding of CsrA leads to the degradation of the target mRNA or prevents ribosomes from binding to the Shine-Dalgarno sequence (55,63,64). CsrA has been reported to negatively regulate all of the known glycogen biosynthetic genes (*glgB*, *glgC*, *glgA*, and *glgS*), which are encoded by the *glgBX*, *glgCAY*, and *glgS* operons, respectively (65).

While CsrB and CsrC do not base-pair with their target mRNA, they serve as a reservoir to control the abundance of free intracellular CsrA. *In vivo* experiment have demonstrated that the cellular glycogen levels and expression of *glg* genes in a *csrB* mutant are closer to those of a *csrB* and *csrC* double mutant compared with the *csrC* mutant (14). Consequently, we used a *csrB* mutant strain to test the effect of downregulating *csrB* and *csr*C expression (**Figure 3**).

Since *glgS* was reported to be positively regulated by CRP (66). Our functional network reveals that CRP, CsrABC and *glg*S form a feed-forward circuit. This circuit resembles an incoherent type 4 FFL. The only difference is that CsrB and CsrC do not interact with the target mRNA of CsrA directly (7,67).

### Characters of the FFLs with sRNA regulators

Most computational simulations and wet-lab experiments have focused on the dynamic patterns in the presence and absence of sRNA. Although mathematical modeling has been applied to analyze the features of different FFLs, proposing that the combination of different promoters generates distinct kinetic functions (37), simulation results indicate that several pairs of FFLs share identical kinetic responses. This suggests that kinetic differences are not the only consideration for organisms selecting different FFLs. Our method enables the systematic identification and characterization of network motifs with sRNAs’ participation, providing sufficient data to evaluate the function of these FFLs. The function and type of a motif can be characterized by fitting the dose-response curve of the target gene in a wild-type strain and an sRNA knockout mutant. To characterize the FFLs with sRNA regulators, we evaluated the effect of FFLs using four metrics: sensitivity (the slope of the dose-response curve); threshold concentration (defined as the concentration of stimulus corresponding to the inflection point of the sigmoid function); and the maximum and minimum expression levels of target genes (**Figure 4 and Table 3**).

**Figure 4.**
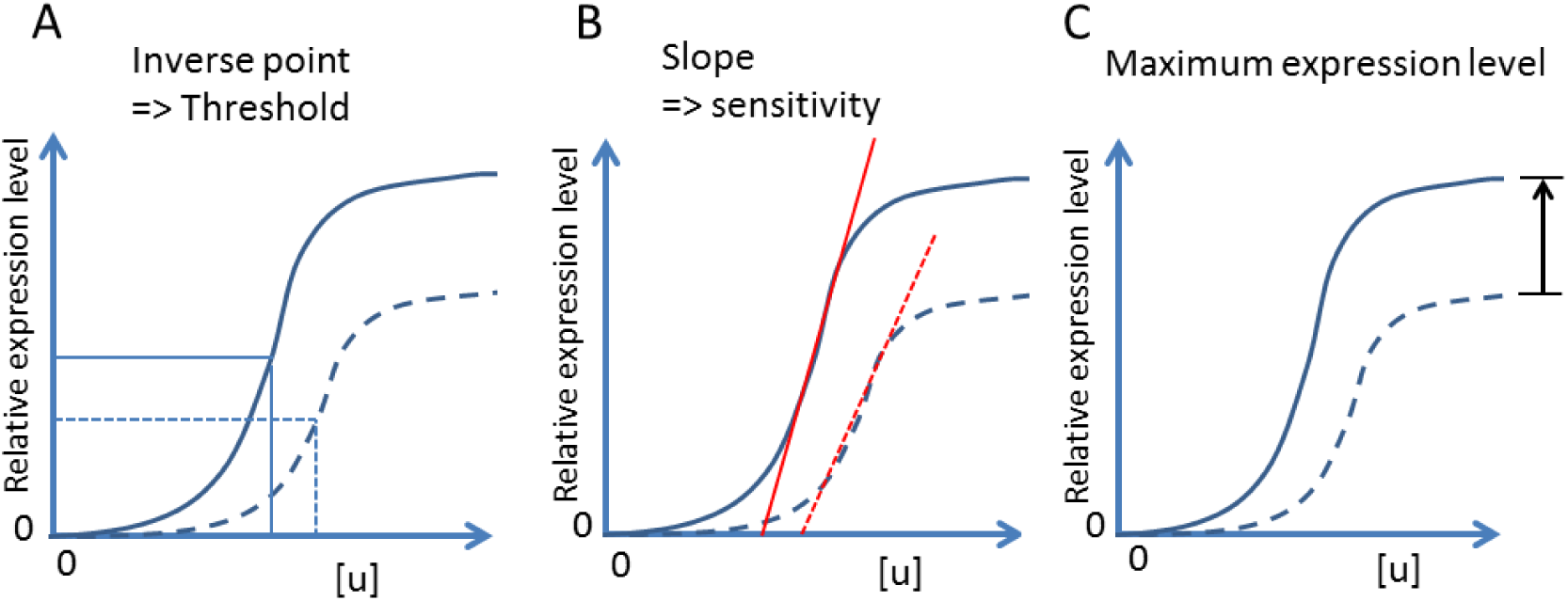
Definitions of characters of a dose-response curve. (A) The threshold concentration is the concentration of stimulus that inducted half the value of maximum expression. (B) The sensitivity is measured by the slope of the signal–response curve on a log–log scale. (C) The difference of maximum expression under different stimulus strength is calculated from the distance of dose-response curve (signal–response curve) in the presence of absent of sRNA.

**Table 3.**
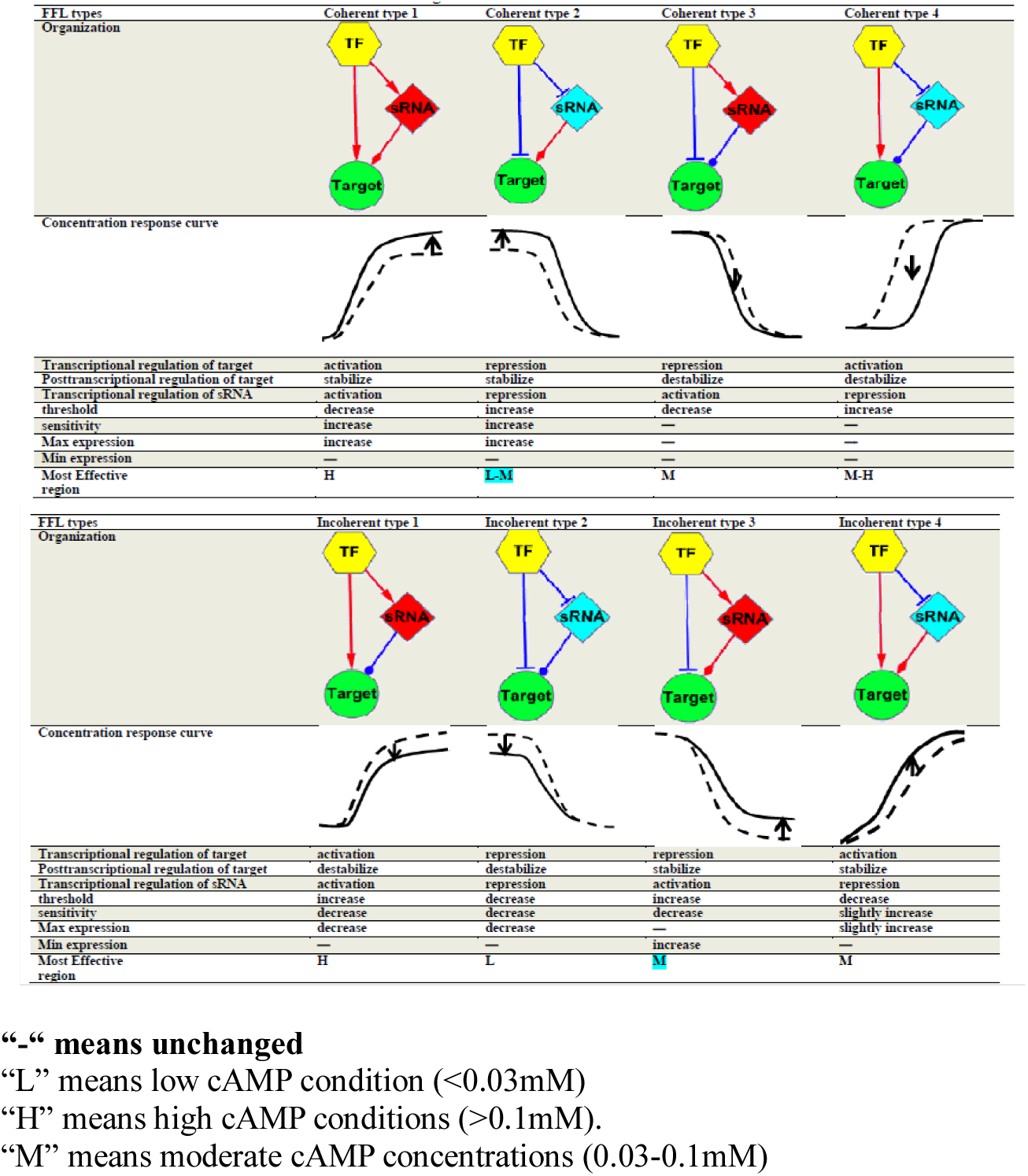
A. Characters of the FFLs with sRNA regulators.

| FFL types | Coherent type 1 | Coherent type 2 | Coherent type 3 | Coherent type 4 |
| --- | --- | --- | --- | --- |
| Organization |  |  |  |  |
| Concentration response curve |  |  |  |  |
| Transcriptional regulation of target | activation | repression | repression | activation |
| Posttranscriptional regulation of target | stabilize | stabilize | destabilize | destabilize |
| Transcriptional regulation of sRNA | activation | repression | activation | repression |
| threshold | decrease | increase | decrease | increase |
| sensitivity | increase | increase | — | — |
| Max expression | increase | increase | — | — |
| Min expression | — | — | — | — |
| Most Effective region | H | L-M | M | M-H |

| FFL types | Incoherent type 1 | Incoherent type 2 | Incoherent type 3 | Incoherent type 4 |
| --- | --- | --- | --- | --- |
| Organization |  |  |  |  |
| Concentration response curve |  |  |  |  |
| Transcriptional regulation of target | activation | repression | repression | activation |
| Posttranscriptional regulation of target | destabilize | destabilize | stabilize | stabilize |
| Transcriptional regulation of sRNA | activation | repression | activation | repression |
| threshold | increase | decrease | increase | decrease |
| sensitivity | decrease | decrease | decrease | slightly increase |
| Max expression | decrease | decrease | — | slightly increase |
| Min expression | — | — | increase | — |
| Most Effective region | H | L | M | M |

“-“ means unchanged “L” means low cAMP condition (<0.03mM) “H” means high cAMP conditions (>0.1mM). “M” means moderate cAMP concentrations (0.03-0.1mM)

The threshold is reduced in FFLs where sRNAs function in parallel with the transcriptional regulation on the target (such as coherent type 1 and 3, and incoherent type 2 and 4). In contrast, FFLs where sRNAs function to antagonize the transcriptional regulation exhibit an increased threshold (such as coherent type 2 and 4, and incoherent type 1 and 3). Regarding sensitivity, coherent type 3 and 4, as well as incoherent type 1, 2, and 3, tend to reduce sensitivity. Conversely, coherent type 1 and 2, and incoherent type 4, all increase sensitivity. Only the incoherent type 3 FFL was found to reduce the basal expression level of its target. Coherent type 1 and 2 FFLs elevate the maximum expression level of the target. In summary, each type of sFFL serves as a distinct response element within the CRP regulon (**Table 3**).

### sRNA-mediated FFLs direct the dynamics of the CRP regulon along with cAMP concentration

The characteristics of sRNA-mediated FFLs summarized in Table 3 not only help to clarify the relationships among factors but also quantitatively explain the observed differential expression levels of the sRNA-target genes between the wild-type and sRNA deletion strains. By investigating the effective regions of different FFLs, we found that they can serve to distinguish different functional gene groups (**Figure 5**). At low cAMP concentration, the regulatory effect of coherent and incoherent type 2 FFLs is most evident. A variety of stress response genes were found to be controlled by coherent and incoherent type 2 FFLs, whose effective region falls within the low cAMP concentration range (< 0.03 mM). For example, the sRNA DicF regulated *FtsZ*, an essential gene for cell division. During cold temperatures, DsrA helps stabilize *rpoS*, which encodes an alternative sigma factor for the general stress response (68), and *hns*, which encodes a histone-like protein that represses the expression of many genes (69). Moreover, other genes, such as *luxS*, which is involved in quorum sensing, and *ompX*, a porin expressed in the outer membrane stress, are expressed when cell density is high (70,71). Furthermore, Gene Ontology (GO) analysis reveals that sFFLs within the same regulatory niches indeed involve closely related biological pathways or cellular activities. These suggests that the topology in an sFFL is subject to evolutionary selection rather than being formed by a random process.

**Figure 5.**
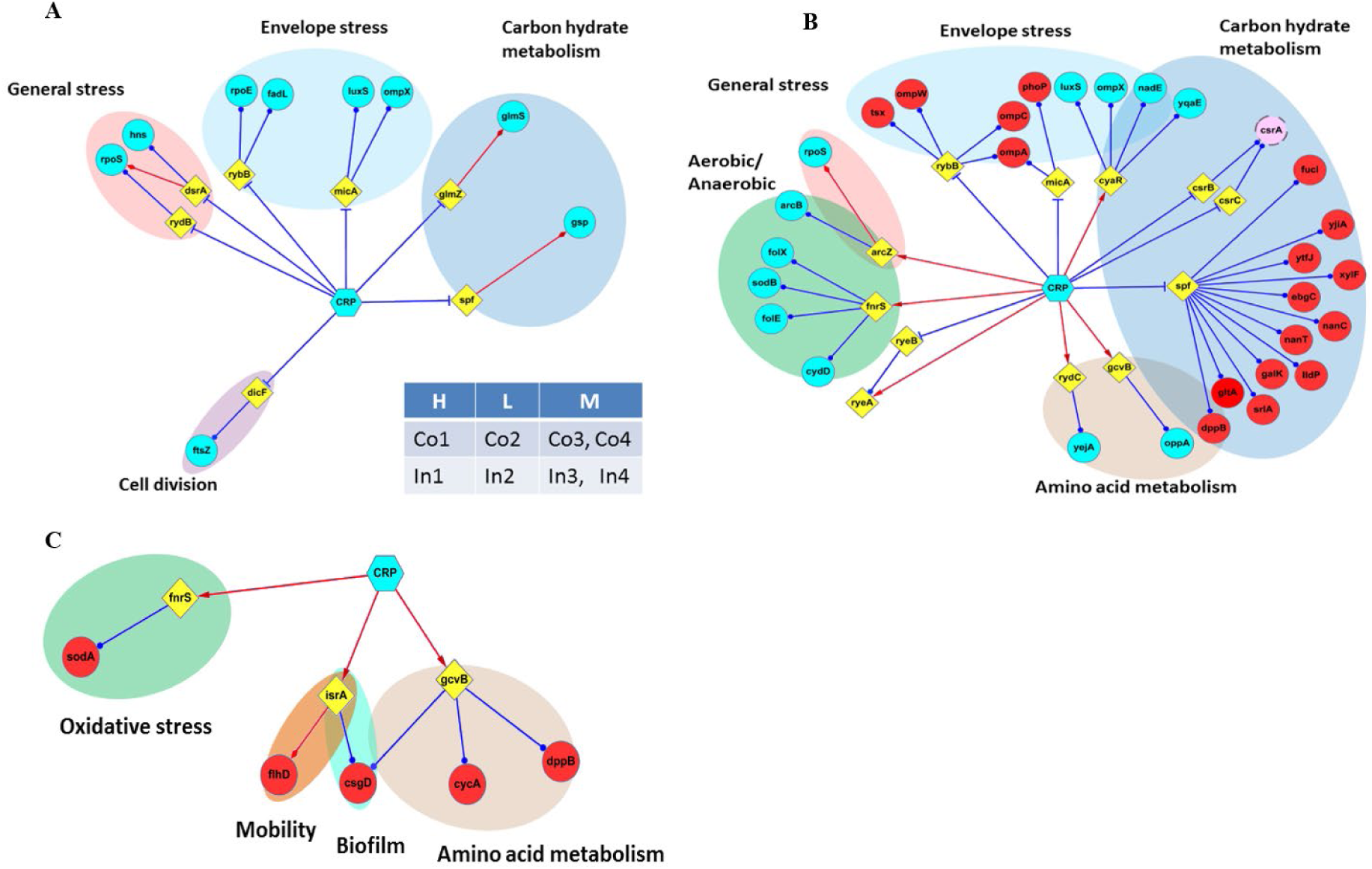
CRP-regulated sRNAs classified by the effective region of each FFL. The effective regions of different FFLs were used to classify the function network of CRP-regulated sRNAs. In this plot, the effective region of each FFLs is defined by the experiential data of selected FFLs (Table S2). The three categories were divided by the cAMP concentration. (A)The first category works under low(L) cAMP condition (<0.03mM) and includes coherent and incoherent type 2 FFLs. (B) The second group encompasses coherent type 3 and 4 along with incoherent type 4. All of these FFLs serve on moderate(M) cAMP concentrations (0.03-0.1mM). (C) The third group composed of coherent type 1, incoherent type 1 and type 3, which are most effective under high(H) cAMP conditions (>0.1mM). Noteworthy is that the effective region of some FFLs can locate to a broad cAMP concentration rage. The abbreviations of prefix, ‘C’ and ‘IC’, present coherent type and incoherent type separately. It seems that genes with similar function tend to be controlled by sFFLs with similar effective region, suggesting the presence of evolution stress in favor of different sRNA regulate FFLs.

## DISCUSSIONS

### Validation of CRP binding sites and prediction reliability

Initially, we selected the threshold value that allows 90% of the training sequences to pass, based on statistical probability. Our EMSA results confirmed that this threshold is reasonable. All 35 candidates with CRP binding sequences on the sRNA promoter region showed distinct retarded bands in the EMSA assay, which have a predicted score greater than 6.0. The promoter fragments of *psrN, ryfB* and *micA*, which have prediction scores near the threshold, showed weak binding affinities with CRP (**Figure 2A**). In contrast, among 21 promoter sequences with a lower cut-off score from 4.9 to 6.0, only two sequences were identified to have weak binding affinity to CRP. Of the 19 out of 21 targets, DNA fragments do not show any clearly shifted band with CRP, except for two DNA fragments of *symR* and *rydC* (**Figure 2B**). These results demonstrate that our scoring method effectively identifies most CRP binding sites in sRNA promoter regions. The difference in binding affinity can be attributed to the specific sequence determinants of the CRP binding site (72). This residue enables CRP to recognize a GTG motif. An ideal CRP binding site encompasses a GTG motif near the 5’ end and a CAC motif near the 3’ end. The predicted binding sites for *ryfB* and *psrD* both contain a GTG motif; however, the latter has a poly-A motif, which likely renders the interaction between the sequence and CRP weak. The binding site on *istR* has a CTG and a CAG, which CRP does not favor. A point mutation study has shown that the interference effect of a point mutation is to replace the conserved GTG with CTC, rendering the binding affinity of CRP to this sequence insignificant (73). The tendency was also distinguished by the sequence having a score higher than 6.0 compared with those scored between 4.9 and 5.9. If a sequence motif is defined as each base pair conserved above 60 percent among a sequence cluster, the motif for all highly scored sequences is TGTGA-6-TCAC, but that of sequences scored between 4.9 and lower than 6.0 is TGA-6-ACT (**Supplementary Figure S3**). Clearly, the sequence motif essential for CRP binding is more conserved in the sequences that pass our threshold value, which is in agreement with biological evidence. Therefore, combining biological information with computer prediction can enhance the accuracy of prediction and aid in interpreting the reliability of the prediction.

### The presence of the CRP binding site alone does not guarantee a dose-response

In this study, we found that while EMSA has confirmed the presence of CRP binding sites in the promoters of some genes, the expression levels of these genes remain unchanged over the test range of cAMP concentrations. For example, in contrast to *ompX* and *luxS*, CyaR is indispensable for CRP-mediated regulation of *nadE* and *yqaE* (**Supplementary Figure S1B**). The cAMP concentration-dependent repression was abolished in the *cyaR* mutant, despite the presence of a CRP binding site in the promoter region. These examples provide a lesson: to determine the presence of an FFL during specific growth conditions, one cannot only consider the regulatory relationships between each factor. However, it does not necessarily mean that CRP cannot control the expression directly. Since other transcription factors may cooperate with CRP to regulate the expression of target genes, one can expect that the function of CRP may depend on the presence of other co-activators and co-repressors. Alternatively, the promoter of these genes may be occupied by other transcription factors during the exponential phase. Indeed, similar results were also reported in a systematic study using ROMA (56). The complex nature of gene regulation may explain this.

In some cases, hetero-cooperation is required for regulated gene expression. For example, the repression effect of CRP on *udp* promoter is dependent on the presence of CytR (74). In other cases, CRP competes with other transcription factors for an overlapping binding site, and thus, the regulatory effect may be masked when other transcription factors dominate the regulatory region (75). A recent study about *fnrS* demonstrated the complexity of sRNA expression under different conditions (49). It was reported that CRP acts as a transcriptional activator for *fnrS* under aerobic growth conditions. Conversely, CRP serves as a repressor when cell growth occurs anaerobically. The ambiguous role of CRP is due to the CRP binding site overlapping with that of FNR, a global transcriptional factor that is only expressed during anaerobic growth. Although both FNR and CRP serve to activate the transcription of *fnrS*, transcriptional activity of the former is about tenfold stronger than the latter. Thus, the binding of CRP reduced the activation of *fnrS*. Moreover, ArcA is found to cooperate with CRP to activate the *fnrS* anaerobic expression in an *fnr* mutant strain. The collection of the above evidence reveals the expression of sRNA is tightly control by many environmental signal inputs. We have demonstrated that the CRP regulation for some sRNAs is more pronounced under specific growth conditions, such as the log phase or low temperature (Figure S4). However, it would be a huge project to detect the concentration response of all sRNAs under any growth conditions that have been reported to stimulate the expression of a particular sRNA. Here, we demonstrate that the cAMP concentration response can be observed in most reported sRNAs in E. coli during cell growth in LB broth under aerobic conditions. Therefore, we believed that most of the regulatory effect can be detected in normal experimental conditions.

### A unique role of sRNAs in modulating the target gene in the network

A recent review article hypothesized that all of the major transcription factors regulated at least one sRNAs (5). Our systematic examination, which extends knowledge about CRP-regulated sRNAs, supports this theory. The major difference between sRNAs and transcriptional factors is that sRNA regulates their targets at the post-transcriptional level. sRNAs do not work with promoters to perform AND- and OR-gates, because sRNA neither targets the promoter nor cooperate with other transcription factors to regulate transcription. Furthermore, sRNA cannot directly activate or repress the expression of its target gene, but rather elongate or shorten the half-life of the target mRNA (37). We therefore believed that FFL, comprising sRNA, must have some specific characteristics that explain the different evolutionary advantage of FFL usage in a regulatory network.

While it is generally recognized that CRP regulates multiple targets in a single-input manner, our new network reveals that CRP cooperates with many sRNAs to regulate a number of genes, which is referred to as a multi-output feed-forward loop (**Figure 5**). More strikingly, while coherent type 1 and incoherent type 1 motifs have been reported to be the dominate motif in the regulatory network and CRP-regulated sub-network *in E. coli*, we only found one coherent type 1 FFL in our network (1,76,77). One possible explanation is that most of the reported sRNAs serve as repressors for target genes, and thus, coherent type 1 FFL is scarce in the sRNA network. This explanation, however, is undermined by the fact that examples of both coherent type 2 and incoherent type 3 FFL can be found in our network. An alternative explanation is that the dominant FFL type may differ in FFLs comprising sRNA and those encompassing transcriptional factors as well.

Admittedly, while our systemic approach identified most CRP-regulated sRNAs, the calculation results are limited by the reported direct targets of sRNA. Thus, there must be some bias that happened because we failed to identify all the target genes of CRP-regulated sRNA. We have also not determined whether this result can be applied to other global transcription factors. However, our workflow will be helpful in finding answers to many further questions: whether the usage of FFLs characterizes the regulatory network of different transcription factors, whether sRNA and transcription factors prefer to participate in some specific FFLs, and whether an organism prefers to use a specific FFL under different conditions. Interestingly, in addition to CRP-regulated sRNA, the dominate FFLs of Fur or Fnr-regulated sRNA are also enriched with coherent type 3 and 4 as well as incoherent type 1, similar to the frequency of the FFLs in *crp*-regulated sRNA (**Supplementary File S2**).

### Functional segmentation of the CRP regulon by FFL effective regions

Most CRP-regulated sRNA functions under moderate cAMP concentration (0.03<X<1mM) (**Figure 3**). Although a reliable estimation of intracellular cAMP concentration is unavailable, it is generally assumed that 1mM extracellular cAMP is sufficient to fully activate the lac operon in the *cyaA* mutant strain(61,78,79) and that a *cyaA tol*C double mutant strain requires only 0.1mM cAMP to reach the peak of expression(61). As a result, general experimental conditions fall within the range of 0.03 to 1 mM extracellular cAMP, where the difference in dose response between the cyaA and cyaA tolC mutant strains is most significant in our dose response curve. This region parallels the effective region of coherent Type 3 and 4, as well as incoherent Type 3 and 4 FFLs (**Figure 5**). Therefore, most of the target genes of CRP-regulated sRNAs fail in this region. Coherent type 3, which serves to reduce the repression threshold, dominates the genes involved in the aerobic/anaerobic transition and amino acid metabolism.

In contrast, coherent type 4 FFLs, which increased the activation threshold, govern carbohydrate metabolism and envelope stress genes. Although no incoherent type 4 FFLs were found in our network, we were surprised to find that the expression profiles of some target genes of the CsrABC system were similar to the simulation results of incoherent type 4 FFLs. Although there were no CRP binding sites on the *csrA* gene, CsrB and CsrC sRNAs, which sequester *csrA* from binding targets, and some target genes of the CsrA protein were found to be regulated by CRP simultaneously. This new finding provides a comprehensive picture: all the sRNAs that respond to carbohydrate metabolism, such as spf, gcvB, cyaR, and csrBC, are regulated by CRP, and almost all of their target genes are regulated by sRNA-mediated FFLs.

Since the transcription of sRNAs correlates with the cAMP concentration, only sRNAs that are positively regulated by CRP function in high cAMP concentration (>1mM), where sRNA-mediated regulation on coherent type 1 and incoherent type 1 is most significant. Coherent type 1 FFLs were only found to regulate *flhD*, whereas incoherent type 1 FFLs could be found to regulate sodA, an oxidative stress gene, dppA, cycA, and *rpoS*, an alternative sigma factor.

In contrast, FFLs with CRP-repressed sRNA typically operate under low cAMP concentrations (<0.03 mM). Most of them involve different stress responses. For example, *rybB* and *micA* are critical regulators in the envelope stress response, and some of their target genes, such as ompX and luxS, are also subject to regulation by another sRNA-mediated FFLs, usually of the coherent type 3 FFLs. Another group includes two regulators of *rpoS,* which encodes the alternative sigma factor S. DsrA is an essential gene that stabilizes the expression of rpoS during cold temperatures (80). In contrast, *rydB* is a destabilizer that limits the expression of *rpoS* when cell culture enters the stationary phase (81). Such regulation may be associated with the fact that intracellular cAMP levels drop sharply after cell growth enters the pre-stationary phase.

### Integration of network motifs diversifies output responses

Investigating the interaction between connected network motifs remains a challenge to systems biology. The participation of different sRNAs and the organization of various network motifs enable the generation of diverse response patterns through a single transcription factor. Our study provides a real-world model to examine previously computational simulation results, and sheds light on the study of complex network motifs. There are seven genes found to be targeted by more than one CRP-regulated sRNA.

From the above analysis, we know that there are three conditions where multiple FFLs regulate a gene. First, the combination of FFLs with different effective regions enables CRP to generate a more complex dose-response curve compared with those genes controlled by a single FFL. Examples can be found in the target of CyaR and Spf. LuxS and *ompX* are regulated by a coherent type 3 FFL involving CyaR, and an incoherent type 2 FFL with MicA. The effective region of the former is located in the moderate cAMP concentration region, whereas the latter is more effective in low cAMP conditions (Figure S7). Similarly, two Spf-regulated genes, *srlA*, which encodes a subunit of the glucitol PTS permease GutABE, and dppB, which encodes a subunit of the dipeptide transporter dppABCDF, are not only regulated by coherent type 4 FFLs with Spf but also controlled by incoherent type 1 FFLs involving *gcvB* and *srlR*, respectively. Interestingly, *srlR* is a CRP-regulated transcription factor rather than an sRNA.

In the second place, a target gene may be regulated by different FFLs under different environmental conditions. The expression of sRNAs that participate in these FFLs is subject to additional regulation other than CRP. One of the most obvious examples is the post-transcriptional regulation of the alternative sigma factor σs, which controls the expression of several genes involved in diverse stress responses, including acid tolerance, the aerobic-anaerobic shift, nitrogen starvation, and entry into the stationary phase. Because of the important role of σ^s^ on gene expression, it is tightly controlled at the levels of transcription, translation, and protein stability (82). To date there are four sRNAs have been reported to regulate the stability of *rpoS* at post post-transcriptional level. DsrA and RprA were demonstrated to positively regulate the transcription of *rpoS* by base-pairing with the sequence hindering the Shine-Dalgarno sequence (83,84). The expression of DsrA peaks when cells are grown in low temperature (25℃) (85), whereas that of RprA is stimulated by increasing the osmolality of the cell. In addition, while OxyS negatively regulates the stability of *rpoS* under oxidative stress, ArcZ is expressed under aerobic conditions and stabilizes the *rpoS* transcripts (48). In this study, cells were grown in LB aerobically at 37℃. Thus, our array only detected a significant concentration response of ArcZ, as well as RydB and DsrA, in a minor population. According to their dose-response, they form incoherent Type 3 and Type 2, as well as coherent Type 2 FFL (**Supplementary Figure S7**). The lacking of *oxyS* demonstrated that not all of the circuits present in the interaction network work simultaneously.

Furthermore, the low abundance of DsrA can be explained by the fact that its promoter activity and turnover are temperature-sensitive (80,85). Certainly, the effect of these FFLs peaks under different conditions and dominates the stability of *rpoS*. The example of *rpoS* provides a lesson: it may be a bias to define the presence of FFL based solely on the records about the relationships between genes in the database, because the dynamic nature of regulatory networks may introduce a layer of control at the transcription or post-transcriptional level.

Finally, we also found some examples where the same type of FFLs work together to regulate the expression of a single gene. An outer membrane porin, *ompA*, is co-regulated by two coherent type 4 secretion factors (sFFLs). Moreover, a special case is found where CRP regulates the expression of glmS, which encodes an enzyme that catalyzes the first step of hexosamine biosynthesis. The regulatory RNA GlmZ is required for stabilizing *glmS* mRNA, but it is itself destabilized by binding to YhbJ, a protein that recruits RNaseE (86). In *E. coli*, another regulatory RNA, GlmY, can compete with GlmZ for binding to YhbJ, thereby sequestering YhbJ from binding to GlmZ (10). In the functional network of CRP-regulated RNA, we found that CRP not only directly inhibits the expression of *glmS* but also represses the expression of GlmZ and GlmY. This regulation is implemented through two connected, coherent Type 2 FFLs. The one includes *glmY*, which increased the stability of *glmZ* under low cAMP conditions, while the other stabilized *glmS* (Figure S7).

These examples demonstrate that the integration of different FFLs prunes the dose response. The combination of the same type of sFFL and preventing sFFLs with different effective dose-response ranges from working together can avoid the formation of a “futile cycle”, where the input transcription factor activates or represses two sRNA simultaneously targeting the same gene but have opposite regulatory effects. A long-standing question: why does more than one sRNA target many genes under some growth conditions? Perhaps the combination of multiple sRNA-mediated FFL has evolved to optimize the concentration response and provide flexibility in taking different strategies in response to different input signals from both the cell itself and its environment. The variation in dose response may imply that the metabolic rate and utilization of peptides are different from those of other carbohydrates (87).

### 3.6 CRP communicates with other global regulators via sRNAs

In addition to modulating the dosage response of CRP, our interaction network also reveals that sRNAs enable CRP to coordinate other global regulators, thereby shaping intracellular conditions. One example is that CRP regulates the expression of rpoS, which encodes sigma factor S, and rpoE, which encodes sigma factor E, through small regulatory RNAs (sRNAs). Under normal growth conditions, sigma 70 is the dominant sigma factor during the log phase. When cells enter the stationary phase or face envelope stress, sigma factor 70 gives way to other alternative sigma factors. We found that many CRP-regulated sRNAs are highly expressed during entry into the stationary phase, and their transcription is dependent on sigma factor S. While CRP has been reported to negatively regulate the expression of rpoS by Regine Hengge-Aronis in 1994, the detailed mechanism remains unknown (82). In addition, the study did not explain why the abundance of *rpoS* was lower in the *cyaA* mutant strain compared to the wild-type strain during the stationary phase, despite more rpoS mRNA accumulating in the *cyaA* mutant during the exponential phase. In this study, our computational search found a CRP binding site 71 bp ahead of the transcriptional start site of *rpoS*. This binding site was further confirmed by EMSA (Figure S6; Table S3). The cAMP concentration response curve of *rpoS* in wild-type and *arcZ* mutant strain defined an incoherent type 3 FFL, where CRP represses the transcription of *rpoS*, but activates the expression of *arcZ*, a stabilizer of *rpoS* mRNA. The collection of previous observations and our new evidence indicates that while the CRP-mediated repression was dominant during the exponential phase, where ArcZ expression remains low, the elevation of ArcZ when cells enter the stationary phase helps to increase the abundance of *rpoS*. Hfq is an important chaperon protein that interacts with trans-interaction sRNAs to facilitate pair complex formation (88). In our previous work, we learned that CRP acts as a repressor for Hfq gene expression. To determine whether the function of Hfq is affected by cAMP, we measured the concentrations of hfq transcripts and protein using real-time PCR and Western blot, respectively. Since Hfq is a small 11 kDa protein that forms stable cyclic homo-hexamers (89), we suggest that the expression level of hfq in the *cyaA tolC* double mutant strain is greater than that in the wild-type strain in log phase, rendering the hfq protein may accumulate to a higher level compared with the wild-type strain. Therefore, although the abundance of *hfq* transcripts decreased in the presence of cAMP, the protein level appears to remain constant when the cells are harvested after treatment with cAMP for 10 minutes (**Supplementary Figure S8**).

Another example of CRP regulating another global regulator via sRNA is found in the Csr post-transcriptional system. In *E. coli*, the Csr system comprises the CsrA protein and two small RNAs, designated CsrB and CsrC (90). The system governs a wide range of physiological processes, including glycogen metabolism, biofilm dynamics, and the *flhDC* flagella regulators.

Finally, we found that CRP regulates the transcription of *rnpB*, which is the RNA component of ribonuclease P (RNAse P). The transcripts of *rnpB* bind to the RnpA protein to form an enzymatic complex, which functions to process tRNA and 4.5S RNA. Explains why CRP-regulated sRNA targets many genes involved in amino acid metabolism, peptide uptake, and the synthesis of coenzymes, such as tetrahydrofolate. It is likely that CRP not only regulates specific genes to meet the requirements of environmental changes, but also prepares the genes in a suitable condition to ensure they function effectively and efficiently.

## CONCLUSIONS

In this study, we presented a systematic methodology for identifying feed-forward network motifs encompassing sRNAs and their target genes co-regulated by the same transcription factor. By leveraging the cAMP-dependent DNA binding affinity of CRP, we established a robust model to investigate the role of sRNAs within various network motifs. Our findings demonstrate that measuring concentration-response curves can not only characterize distinct feed-forward loops (FFLs) but also uncover the presence of unknown regulators within a pathway. The integration of previous research on FFL kinetics with our dose-response data reveals that each FFL topology functions as a specific control module. This implies that the expression patterns of cellular genes reflect an adaptive program employed by the cell to cope with fluctuating growth conditions. Unlike static electronic circuits, the topology and interaction strength within biological networks exhibit plasticity, easily altered by modulating the concentration of repressors, cooperators, or regulators. A deeper insight into this complete regulatory network enables us to evaluate the distinct evolutionary advantages of using sRNAs versus transcription factors. Furthermore, understanding how different FFLs coordinate will facilitate the design of simplified artificial biological circuits and elucidate how organisms make specific decisions in response to intracellular and environmental signals. Ultimately, this research paves the way for applying network motif concepts to understand complex biological phenomena, ranging from memory and asymmetric cell division to the mechanisms underlying complex diseases.

## Acknowledgements

We want to thank Dr. Steve Busby of the School of Biosciences at the University of Birmingham for donating the pRW50, pDU9, and pDCRP plasmids.

## MATERIALS AND METHODS

Detail methods are described in **Extended Experimental Procedures.**

### Bacterial strains and growth conditions

All mutant strains used in this study were *E. coli* BW25113 derivatives and generated from the Keio collection system, which was kindly provided by the National Institute of Genetics of Japan (Baba et al., 2006). The features of the mutants are listed in Supplementary Table S4. For microarray and Real-time PCR experiments performed to validate array data, wild-type strain BW25113 and its *crp* deletion derivative strain were aerobically grown in 250-ml shakes flasks supplied with 25ml Luria-Bertani (LB) until log phase (OD600 reached 0.4), and then harvest for RNA isolation. For cAMP dose-response experiments, cell were growth in minimum medium supplied with 0.4% glucose until log phase, and then spun down and suspended in fresh minimum medium supplied with 0.4% glycerol and different concentration of cAMP (0, 0.01, 0.03, 0.1, 0.3, 1, 3 and 10mM). Cells were incubated with cAMP contained medium for 10 minutes, and then harvest for RNA isolation.

### Uncover CRP-regulated sRNAs *CRP binding site Prediction*

A CRP-regulated gene has at least one CRP binding site on promoter and shows differential expression between *crp* mutant strain and wild type strain. A CRP binding sites prediction model employed to scanning the promoter of sRNAs. In brief, the sequences of 165 non-redundant, experimental confirmed CRP sites were collected to build a hidden Markov model profile and a weight matrix specified CRP binding sites in *E. coli* by using *hmmbuild* in the HMMER package 3.0 programs (Finn et al., 2011) and *matrix2lib* in the program MATCH (Kel et al., 2003). The default cut-off value is defined as 6.0, which enables 90% of the cis-elements to pass the threshold. **Detail information about this program is described in Extended Experimental Procedures.**

#### Electrophoresis mobility shift assay validates Predicted binding sites

Predicted CRP binding sites scored above 4.9 were further confirmed by electrophoresis mobility shift assay (EMSA). The general protocol was the same as described previously (Lin et al., 2011). In brief, DNA oligonucleotides were generated by PCR using BW25113 genomic DNA as template with primers listed in Supplementary Table S5. CRP protein (His-tag) was prepared by transformed *E. coli* BL21 (DE3) with pQE30-crp plasmid and isolated using His-tag column system (Qiagene, Hilden, Germany). DNA oligonucleotides (20nM) were incubated various concentrations of CRP (0, 0.05, 0.1, 0.2, 0.4, 0.8µM) in a buffer solution containing cAMP (200μM), and Tris–HCl (20mM, pH 8.0), MgCl_2_ (10mM), EDTA (0.1mM), and KCl (100mM) at 37 °C for 30 minutes to allow binding. Then, the mixture were mixed with cyber green loading dye (GeneDireX, Las Vegas, NV, USA) and separated by non-denaturing polyacrylamide gel (6%). The bands were visualized by exposure to UV light. The results show score 6.0 is a threshold value of virtual CRP binding with a false positive rate and false negative rate were 0% and 3%, respectively.

#### RNA Extraction, Microarray Processing, and Data Analysis

Total RNA was extracted using the TRIzol® reagent (Invitrogen, Carlsbad, CA) and treated with RNase-free DNase I (Promega, Madison, WI). The quality of RNA was determined by spectrophotometry and gel electrophoresis. Only samples have A260/A280 ratios between 2.0 and 2.1 and yield distinct 23 S and 16 S bands were further used. RNA was converted into cDNA incorporating amino-ally dUTP by using SuperScript III Reverse Transcriptase (Invitrogen). After RNA template was removed by adding NaOH (100mM) and EDTA (25mM), the amino-allyl cDNA was purified by QIAquick Nucleotide Removal kit (QIAGEN) and then converted to Cy5 labeled cDNA (GE, PA25001). The differential expression of sRNA between the WT and *cr*p mutant strains was measured by single-fluorescence microarray hybridization experiments. We designed an *Escherichia coli* 0.155 K Custom Small RNA Array (GEO platform GPL10535). Each array assays were repeated six times. Before hybridization, Cy5 labeled cDNA was mixed with hybridization buffer (Phalanx), heated to 95℃ for 5 minutes, and then kept at 65℃. The proceed working solution was infused into pre-warmed assembled chip and hybridized at 42℃ for 16 hours. After hybridization, the probed array was dissembled and washed by Buffer I (2×SSC, 0.2% SDS), following Buffer II (2×SSC) at hybridization temperature for 5 min, and Buffer II at room temperature for another 5 min. Next, the probed chips were rinsed with Buffer III (0.2×SSC), and then dried by centrifuged. Finally, the chip was scanned at 635 μm wave length by GenePix 4000B array scanner (AXON), and image analysis was captured and analyzed by GenePix Pro 4.0. The expression of four stable-expression genes (*polA*, *motB*, *yihA* and *serW*) were used to normalize all array intensities. The difference of expression level greater than one-fold and its P-value bellows 0.05 is considered significantly.

#### Real-time PCR

The total RNA (3μg) was pretreated with DNaseI, and then reverse transcribed into cDNA by using SuperscriptIII (Invitrogen). The primers for real time-PCR were designed with Primer Express (Applied Biosystems) and listed in Supplementary Table S6. The reaction mixture (25ul) contained SYBR Premix ExTag (1x)(Takara, Tokyo, Japan), ROX^TM^ Reference dye (1x)(Takara), forward/reverse primers (0.2uM), and properly diluted cDNA template (2ul). All of the reactions were performed by ABI PRISM^®^ 7000 (Applied Biosystems, Foster City, CA). The program and was set as the manufacturer’s recommendation. The related gene expression was calculated by comparative CT method.

#### Northern-Blotting assay

Northern blot was applied to quantity the abundance of sRNA species that only expressed some specific conditions. The probes were generated by using PCR DIG probe synthesis kit (Roche) to incorporate DIG-dUTP during PCR amplification. The RNA samples (10μg) and marker (Ambion, AM7145) were electrophoresed in polyacrylamide gel (8%) with urea (7M), transferred to Millipore charged nylon membrane by Trans-blot semi-dry transfer cell (Bio-Red, Richmond, CA), and UV-cross linked for three times. Before hybridization the membrane was pre-hybridized using FASTHyb-Hybridization solution (Biochain, Hayward, CA) at 65℃ for 30 min, and then the membrane was hybridized with specific DIG-labeled probe in the same hybridization solution at 65℃for one hour. All of the following procedure, including washing, blocking, immunological detection, were performed by using a commercial system (DIG Wash and Block buffer Set, Roche). Finally, the subtract CSPD (Roche) was conversed to luminescence by alkaline phosphate and became detectable in X-ray film (GE Healthcare, Buckinghamshire, UK).

### Validated CRP regulation on sRNA target genes to identify sFFLs

The target genes of sRNA were found by literature review, and all of the reference were listed in a Supplement File S2. The same computational process used to identify CRP-regulated sRNAs was also applied to predicted CRP binding sites on sRNA target genes. Except reported CRP-regulated genes, the differential expression of all target genes of CRP-regulated sRNAs was quantified by Real-time PCR. A target gene with predicted CRP binding sites scored above 6.0 and significant differential expression in *crp* deletion strain is consider to be regulated by CRP directly.

### Quantity Dose-response of FFLs

In cAMP-dose-response experiments, *tolC*, *cyaA* double mutant was used as wild-type strain. Additional sRNA deletion mutant was introduced to wild-type strain by P1 transduction. The general protocol of cAMP dose-response follows the previous reported method with some modification(Hantke et al., 2011). Activated overnight culture was transferred to fresh casamino acid-enriched M9 containing glycerol (0.4%) as carbon source. Cell cultures were pretreated with cAMP at various concentrations (0, 0.01, 0.03, 0.1, 0.3, 1, 3 and 10 mM) for 10 minutes, before harvesting and RNA isolation. The relative expression levels of sRNA or its target gene were determined by Real-time PCR as previously described to quantify differential expression. The Ct of a gene expressed in the absence of cAMP was defined as the standard for calculating the relative expression level under different cAMP dosage.

## Supplemental Material and methods

### Identification of CRP binding sites on sRNAs’ promoter regions

The major challenge to identify the Feedforward loops with sRNAs is that the transcriptional regulation of most sRNAs remains unknown. We therefore developed a systemic approach to investigate the CRP-regulated sRNAs. The computational and experimental approaches are outlined in **Figure M1**. The first step is to build a mathematical model of CRP binding sites based on experimental confirmed CRP binding sites, which is called cis-elements. There are 311 CRP cis-elements in EcoCyc (http://ecocyc.org/) (Keseler et al., 2011). To reduce the effect from the error of data source, there we only considered the elements have been verified by experiments, such as “Binding of cellular extracts”, “Binding of purified proteins” or “Site mutation”. The location of the reported CRP-binding sites were revised by ClustalW (Larkin et al., 2007). Based on these criteria, we removed 54 elements that are redundant or have no positional information, and 92 elements that have no experimental support. Finally, a set of 165 CRP cis-elements was aligned with ClustalW (Fukami-Kobayashi and Saito, 2002) and used to construct the model. A hidden Markov model profile and a weight matrix were built from the reported CRP-binding sites by using *hmmbuild* in the HMMER package 3.0 programs (Finn et al., 2011) and *matrix2lib* in the program MATCH (Kel et al., 2003). In order to filter significant matches out of the high amount of possible matches, the default cut-off value is defined as 6.0, which enables 90% of the cis-elements to pass the threshold. After deciding the cutoff score, the model is used to identify the regulatory sites within the promoter regions (from −500 to +100 base pairs relative to transcriptional start position of sRNAs). Giving the palindromic nature of CRP binding sites, the width of scanning window was defined as 22 base-pairs and scanning from the 5’ end of both strands to score each predicted binding site (Gunasekera et al., 1992). Moreover, if adjacent CRP binding sites over lapping with each other, they would be merged and only the one that has the best score will be defined as a positive site. Through computational screening, there are 51 sites on the promoter of 35 sRNAs. All prediction results and CRP binding sites information were presented on the web site (http://140.113.239.240/~aliken/CRP2sRNA/) (**Figure M2)**.

### Interaction between cAMP-CRP and sRNAs promoter

The computational promoter analysis revealed 51 CRP binding sites of 35sRNAs. The CRP binding ability of all candidate sequences on these sRNAs’ promoter were provided by EMSA to validate the binding ability of CRP to candidate sequence on sRNAs promoter region (**Figure 2**). Amplified DNA fragment was[γ-^32^P] dATP end-labeled and incubated with purified CRP and cAMP (200µM) and then electrophoresed on 6% PAGEs. For 33 of the 36 DNA targets with satisfied cut-off score, clear retarded bands appeared, when the DNA fragments incubated with purified CRP (**Figure 2A**). Furthermore, addition of purified CRP resulted in no retardation in the DNA fragment of *psrN*, *ryfB* and *micA* promoters, which contain predicted binding site with a score near our threshold. On the other hand, among 21 promoter sequences with lower cut-off score from 4.90 to 5.99, only 2 sequences were identified to have weak binding affinity to CRP. The 19 of 21 targets DNA fragment doesn’t show any clearly shifted band with CRP except two DNA fragments of *symR* and *rydC* (**Figure 2B**). These results proved that our threshold score is suitable to identify most CRP binding sites in sRNA promoter regions.

**Figure M1.**
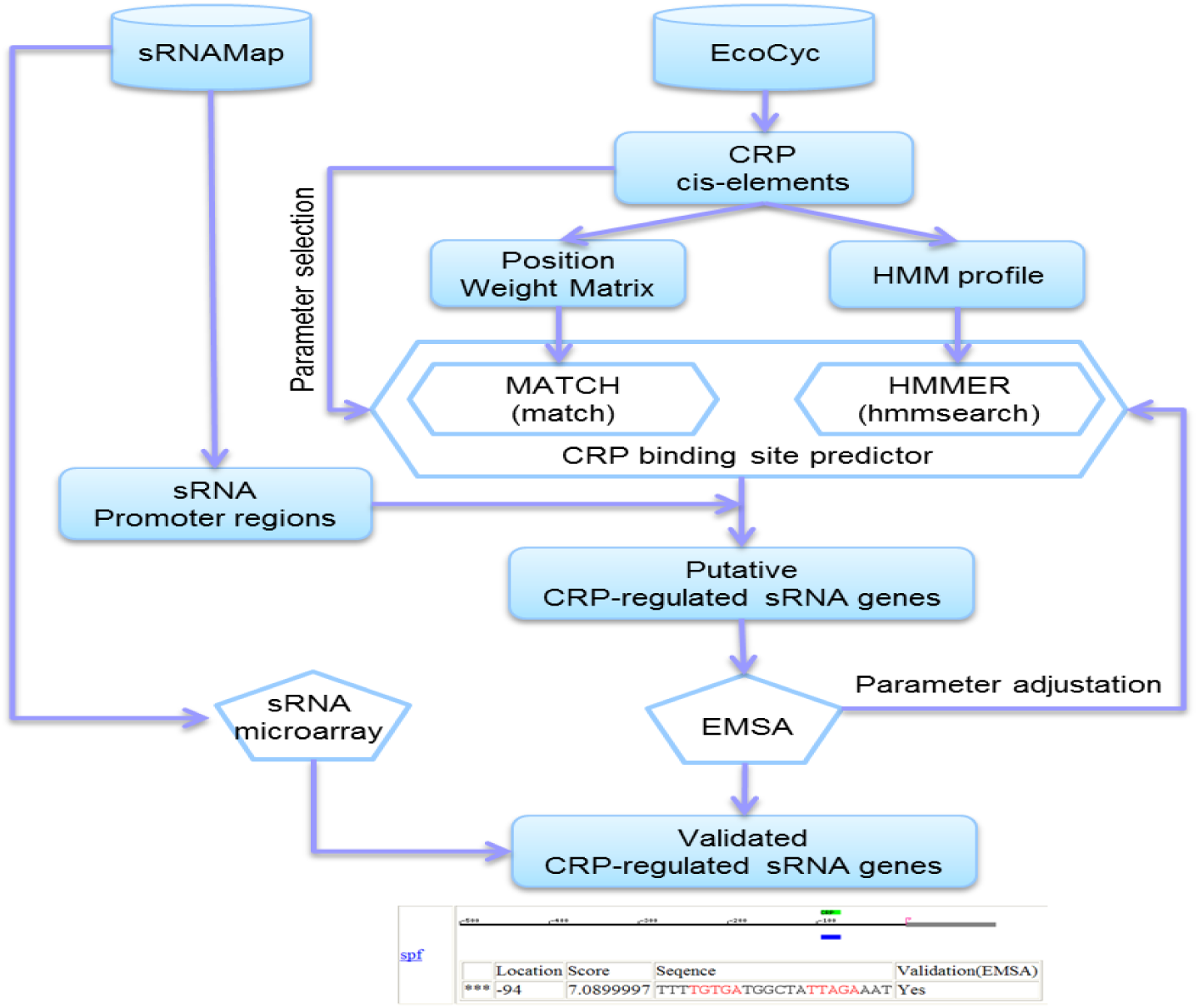
The flowchart of the computational and experimental steps used for identification of CRP-regulated sRNA genes. The strategy involves the following steps: identification of CRP binding sites on sRNAs promoter region, in vitro validation of cAMP-CRP complex binds, and expression analysis of CRP-regulated sRNA genes.

**Figure M2.**
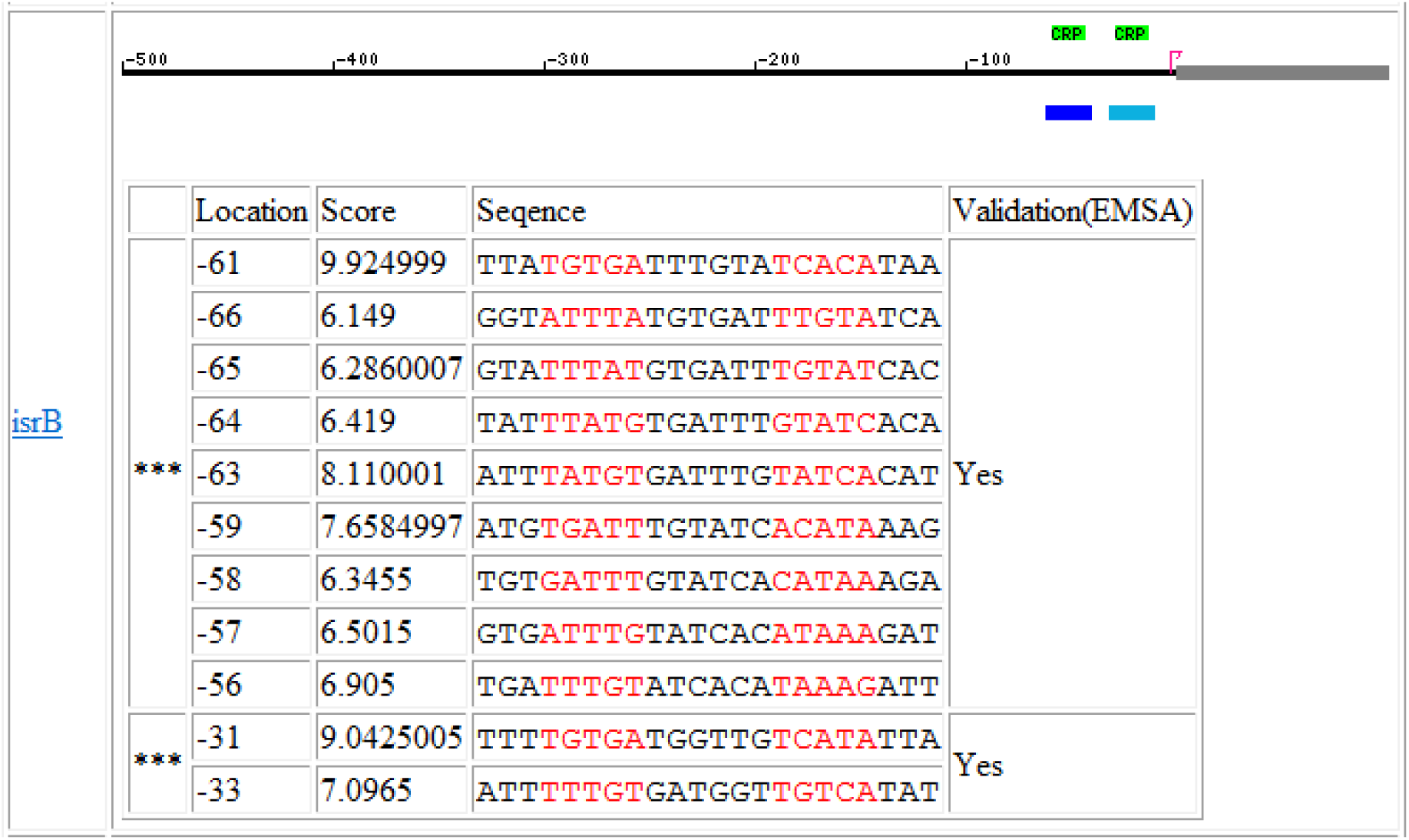
The brief display of the promoter region of CRP-regulated sRNAs. The result web page graphically reveals the location of the identified CRP binding site (blue rectangle: dark blue represents the site have maximum score) and the TSS of sRNAs (red arrow), and the known regulatory information of the sRNAs (green and red rectangle: green represents transcription activation and red represents transcription repression). At the same time, we use text giving information such as the location, score, sequence (core regions are colored red) and status of validation. If the sites cover each other, they would be merged (black star). In the validation status, “Yes” and “Concentration dependent” represent high-affinity CRP binding sites and low-affinity CRP binding sites in the EMSA reaction, respective.

### Focused array analysis of CRP-dependent sRNAs

In order to identify the CRP regulon of sRNAs in *E. coli*, the RNA expression levels in wild-type cells BW25113 and *crp* mutant was examined by single-fluorescence microarray hybridization experiments. An overview of the sRNA focused array results is presented in Figure 2. Microarray. To investigate of CRP-dependent sRNAs in *E. coli*, *Escherichia coli* 0.155 K Custom Small RNA Array (GEO platform GPL10535) was designed. The sequences of the probes were designed by Array Designer 4.25 (PREMIER) (Supplementary Table S7). The probes are synthesized oligonucleotides of 50-mer in length, except that for *dsrA* and *sokA* are in 37 and 30-mer, respectively (Biosearch, CA, USA). There are four groups of probes: 5 exogenous genes from *Arabidopsis thaliana*, 17 stable-expressing genes as internal control, 3 random sequences as negative control and 81 genes coding for sRNAs. There is an array of intensity control generated by spotting serious diluted mixture of 17 stable-expression genes. These probes are spotted as 110μmin diameter onto glass slides using a proprietary non-contact printing method (Phalanx, Taiwan; See http://www.phalanxbiotech.com/ for updated protocols). The detection area was duplicated on a single array, enabling performance two independent assays at the same time. In each area, there are nine independent probes for the same target. Besides, Cy3-labeled position control is located at the surrounding of blocks to facilitate orientation of blocks. By single-fluorescence microarray hybridization experiments, the RNA levels of each sRNA gene were compared between the WT and *cr*p mutant strains. Each array assays were repeated six times.

Among the 82 sRNA genes in *E. coli* K12 strain, the expression level of 35 (43%) sRNAs was greater than 1.5-fold in *crp* knockout strains (*P*-value <0.05). Among them, 11 sRNAs up-regulated and 9 sRNAs down-regulated by *crp* mutant in customized sRNA focus array expression profiles. In addition, 7 CRP regulated sRNAs fail to detection by microarray were proved by Northern Blots assay(**Figure S4**). All of above, in 27 CRP dependent sRNAs, we successfully found 24 novel CRP dependent sRNAs. The three known CRP regulated sRNAs, *spf*, *cyaR* and *fnrS* genes were also identified in our experiments, these confirms that our study were reasonable. We also confirmed the tendency on focused array by Real-time PCR. 6 sRNA genes exhibiting *crp*-dependent regulation were selected for Real-time PCR (**Figure S9**). Overall, we serve to define and extend our understanding of the CRP regulon of sRNAs and revealed novel CRP regulated sRNAs gene regulatory network in *E. coli*.

## Supplemental Figures and Tables

**Supplemental Figure S1.**
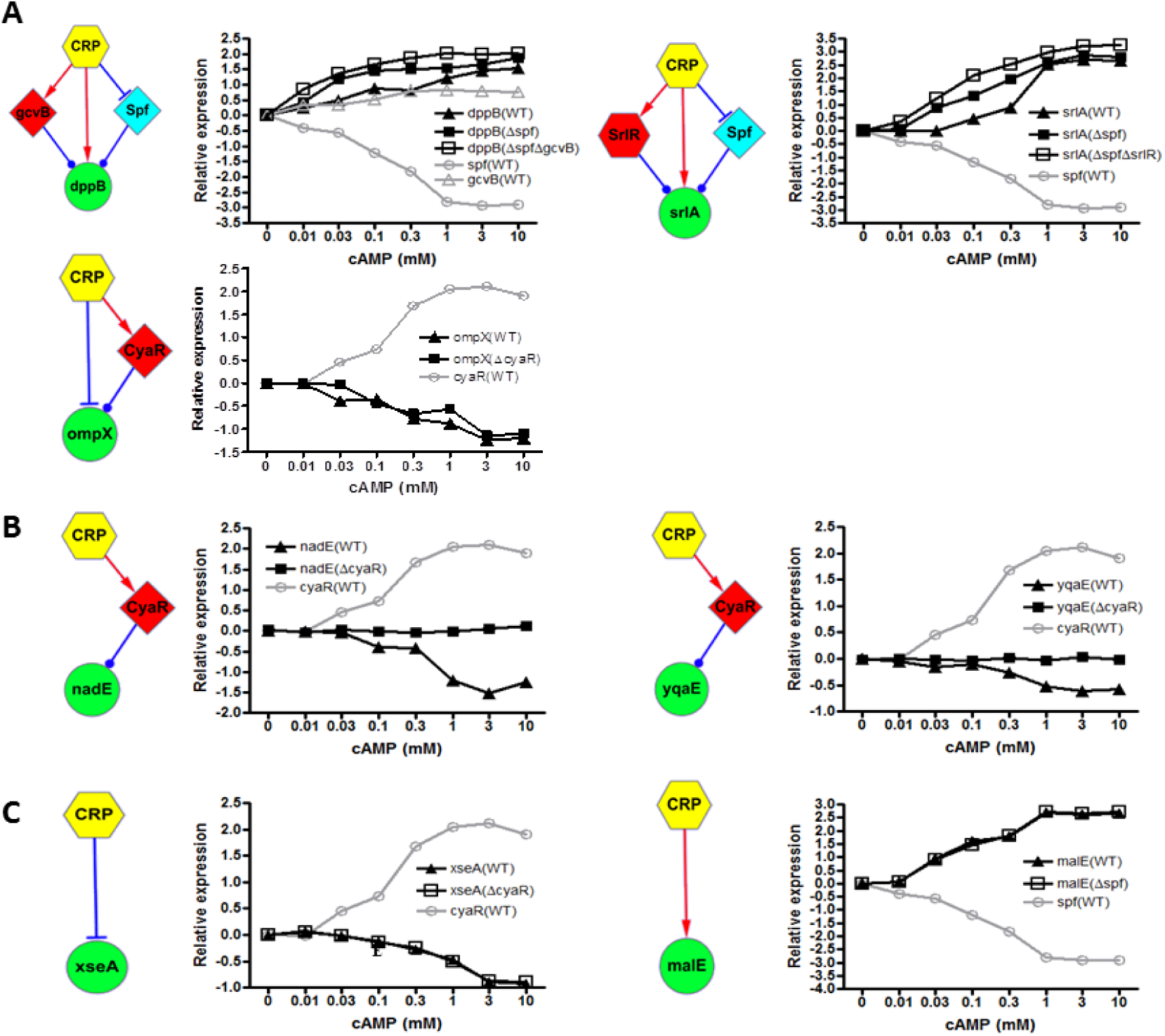
(A)The cAMP dose-response of three sFFLs targeting *dppB*, *isrA*, and *ompX* are illustrated in a log-log scale. (B)The cAMP dose-responses of two sRNA-mediate indirect regulations and(C) two direct CRP-mediated regulations serve as positive and negative control, respectively, for CRP-mediated regulation. The *E. coli* BW25113 derivate Δ*cyaA* Δ*tolC* strain was referred to as the “wild-type” strain (WT) in this experiment, and all of the mutant strains were derivatives of this strain. The response curves of sRNA are draw in gray lines, whereas those of sRNA target genes are illustrated in black lines. The abundance of sRNA (○) as well as target genes in wild-type (▴) and sRNA mutant strains (▪) was determined by real-time PCR. Transcription factor, sRNA, and sRNA target genes are depicted using Hexagonal, diamond, and round symbols, respectively. Noteworthy is that *srlA* is co-regulated by a sFFL and a transcriptional FFL regulated by CRP and *srlR*, a transcription factor. Transcription factor, sRNA, and sRNA target genes are depicted by Hexagonal, diamond, and round, respectively. Noteworthy is that srlA is coregulated by a sFFL and a transcriptinoal FFL regulated by CRP as well as srlR, a transcription factor.

**Supplemental Figure S3.**
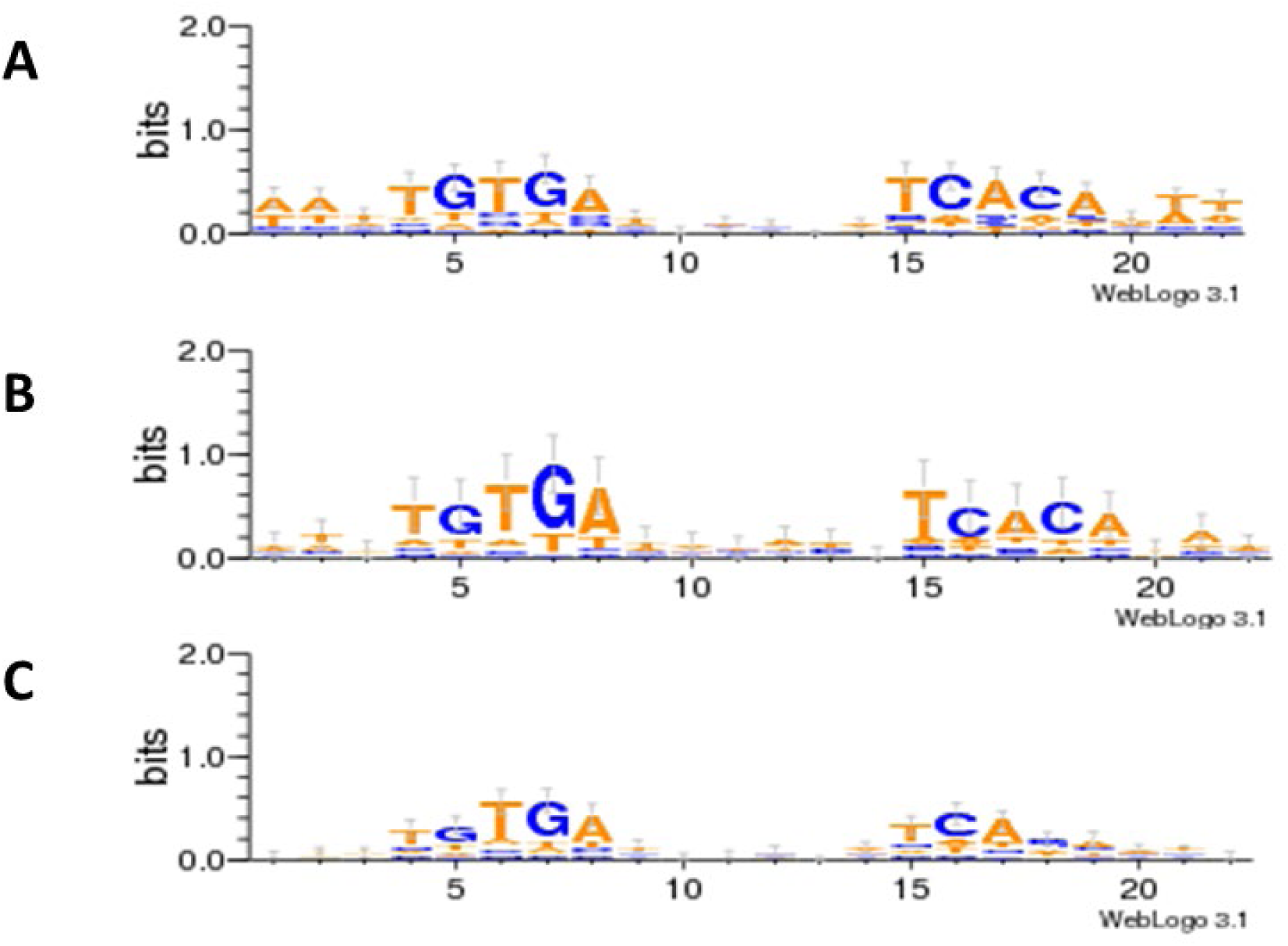
Conservation of predicted CRP binding sites. The sequence logos were generated by Weblogo program(Crooks et al., 2004). (A)The top panel illustrated the conservation of CRP binding site that used to build the CRP binding site prediction model. (B) The middle panel illustrates the conservation of predicted CRP binding sites scored above 6.0, (C) whereas the lower presents the conservation of predicted binding sites scored between 6.0 and 4.9.

**Supplemental Figure S4.**
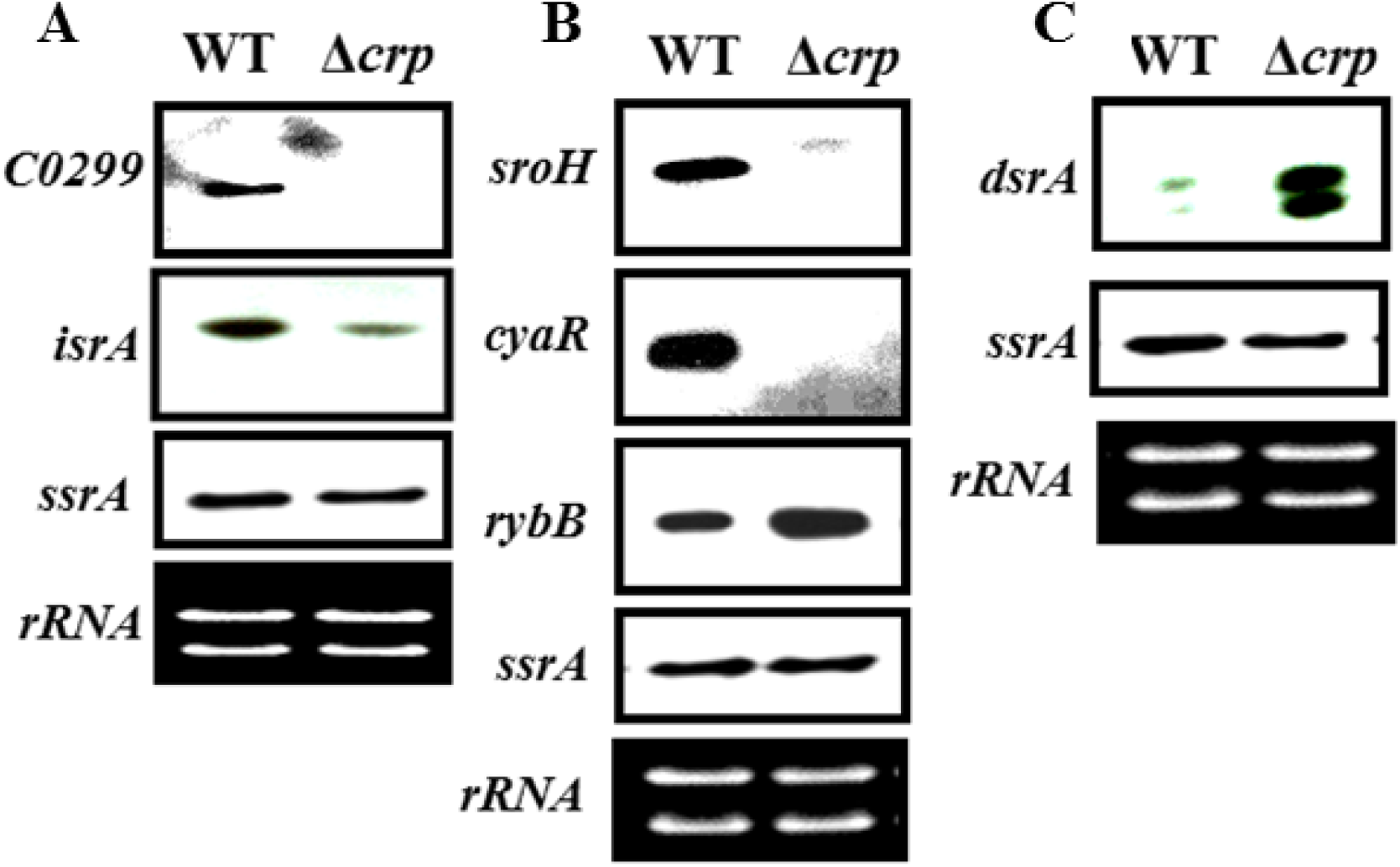
Differential expression of CRP-regulated sRNAs under specific growth conditions. The expressions of six sRNAs fail to be observed on the sRNA focus array in exponential phase. The total RNA extract from wild-type and *crp* mutant strain were analyzed by Northern blot. (A) The expression of *C0299* and *isrA* could be detected in exponential phase, indicting the probe on array may be malfunctioned. The differential expression of other four sRNAs could be detected under specific growth conditions. (B) The abundances of *sroH*, *cyaR* and *rybB* augment dramatically when entering stationary phase. (C) The expression of *dsrA* become detectable when growing at 30℃. The expression of *ssrA* is independent of CRP, and thus *ssrA* serves as an internal control, and the 16 rRNA was used as a loading control.

**Supplemental Figure S6.**
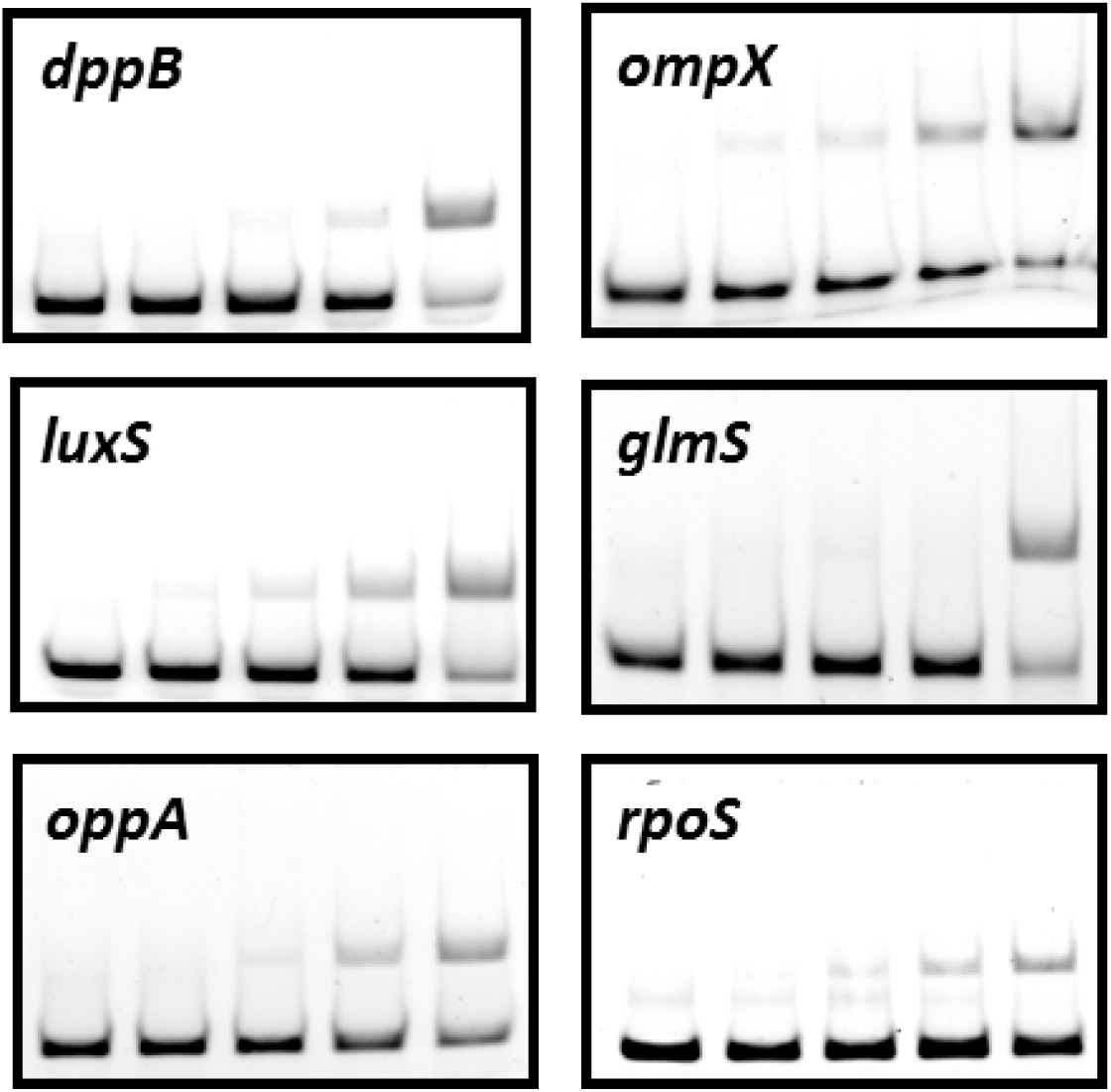
EMSA results of novel CRP target genes. In this study, we predicted some novel CRP binding sites on the promoter regions of some important genes. The CRP binding ability was further confirmed by EMSA. Synthesized oligonucleotides were incubated with cAMP (200μM) and CRP at various concentrations (from 0 to 200 nM) in the buffer.

**Supplemental Figure S7.**
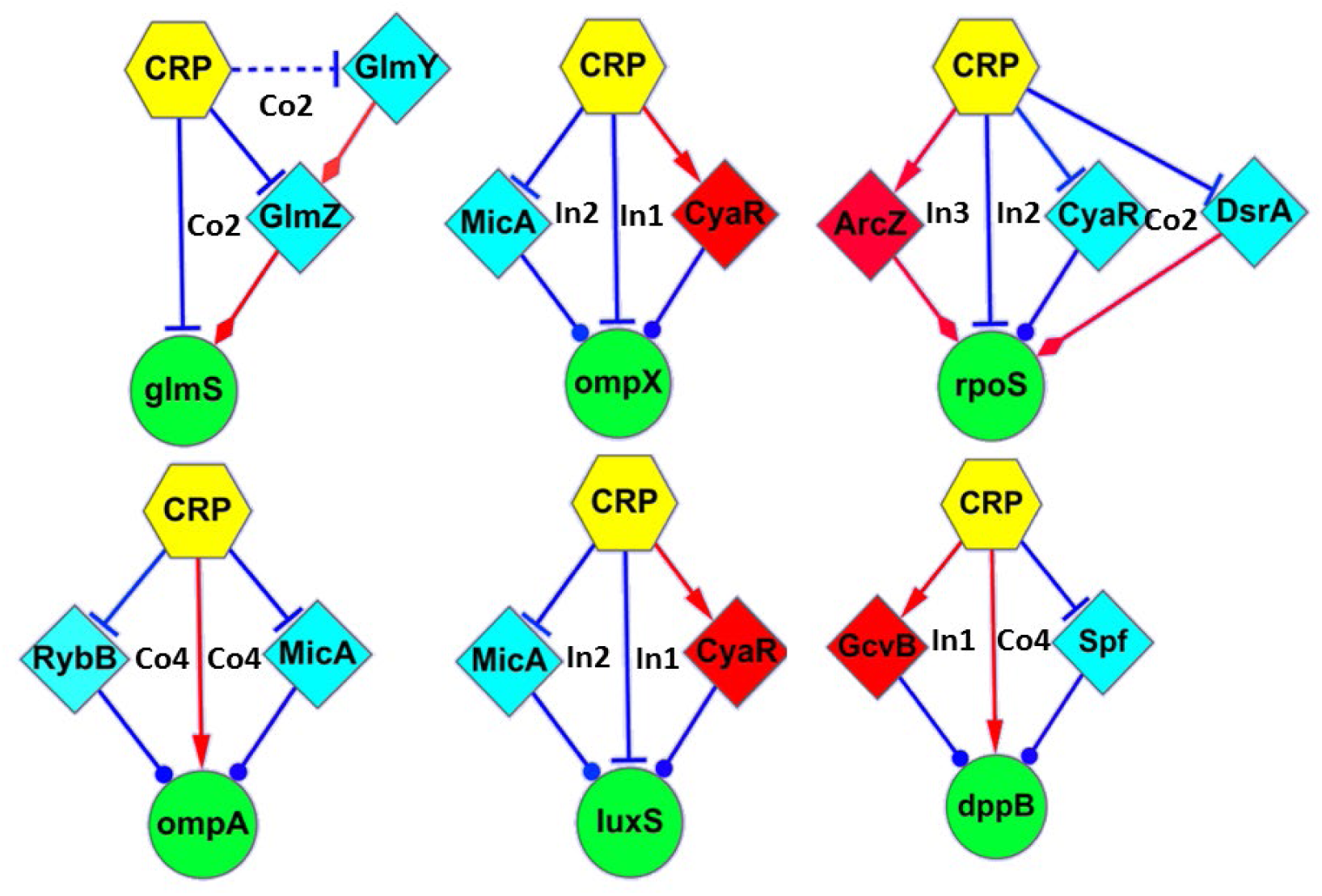
Genes that regulated by multiple sFFLs. The plot summarized all of the genes (or operons) that regulated by multiple sFFLs in CRP regulon. Some of the sFFLs work simultaneously (like MicA and CyaR mediated sFFLs), whereas other function under distinct growth condition. For eammple, DsrA-mediated sFFLs only works under growing at 30℃, because the expression of DsrA is temperature sensitive(Sledjeski et al., 1996)

**Supplemental Figure S8.**
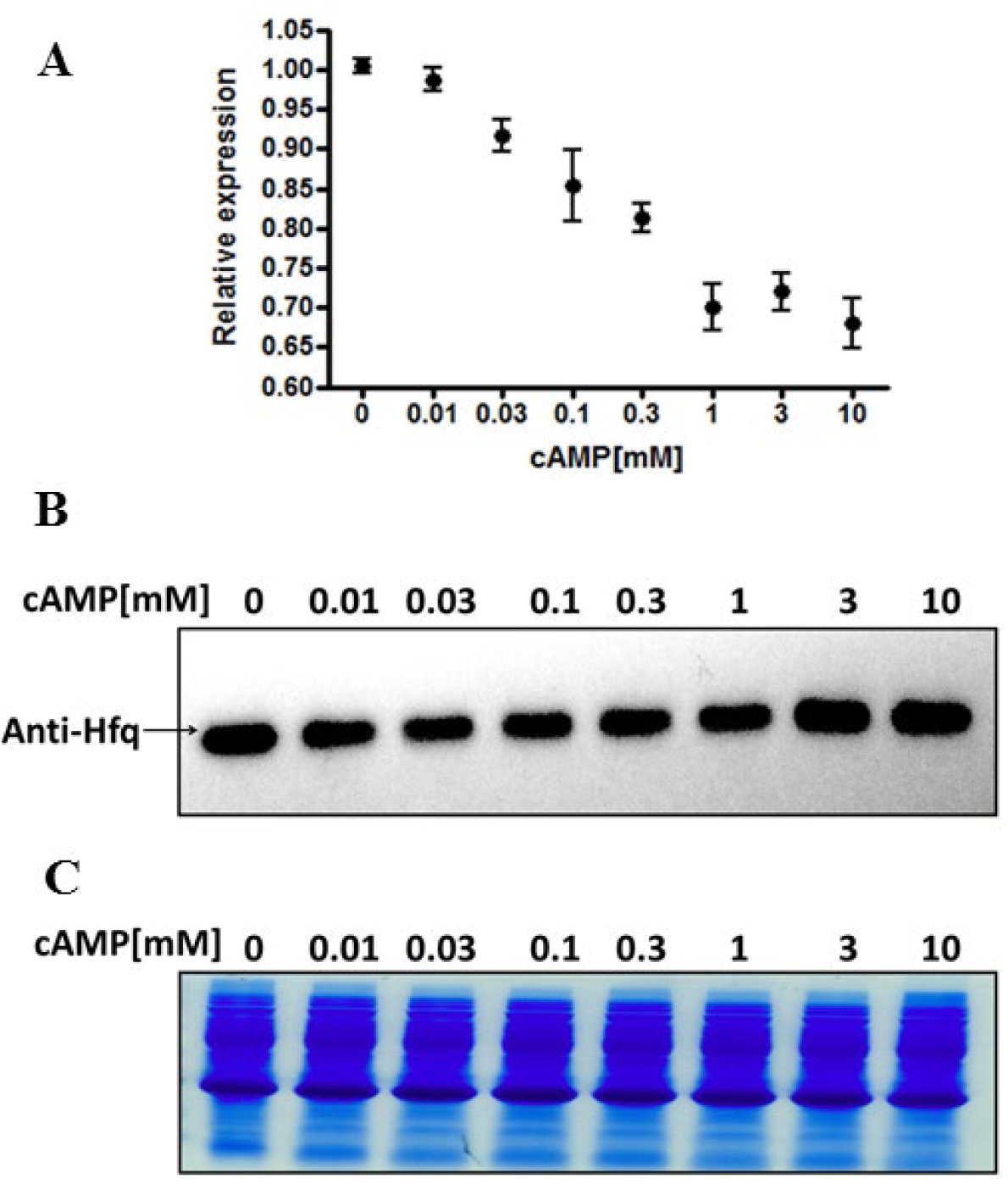
cAMP concentration affect *hfq* mRNA expression level but not protein concentration. (a) The real-time PCR and western blot assay were conducted in Δ*cyaA* Δ *tolC* double mutant strain. The real-time PCR results indicated that cAMP reduces the *hfq* mRNA expression level. This is in agreement with the results in previously study (Lin et al., 2011). (b)The western blots with *hfq* antibody shows that there were no significantly different between Hfq protein expression levels under different extracellular cAMP levels that is used in dose-response experiments. The lower panel indicated the total cellular protein level when cells were growing in different cAMP concentrations.

**Supplemental Figure S9.**
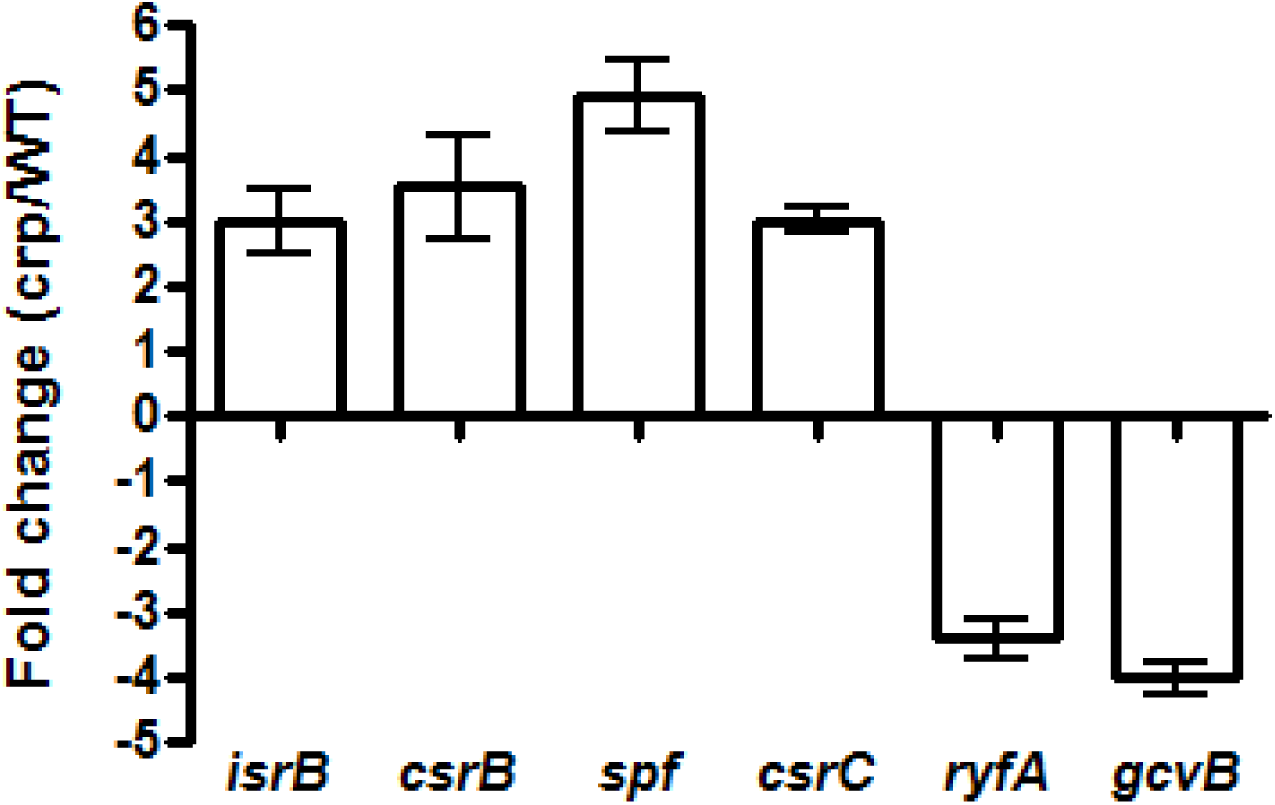
Real-time PCR confirmed CRP-mediated regulation on sRNAs. To confirmed the differential expressions of sRNAs that observed by microarray, the differential expressions of selected sRNA were analyzed by real-time PCR. Comparing the expression levels of sRNAs in *E. coil* BW25113 wild-type strain and its derivative CRP mutant strain, the expression of *isrB*, *csrB*, csrC, and *spf* increased in CRP mutant strain, whereas those of *ryfA* and *gcvB* decreased in CRP mutant strain. These results confirmed the differential measured by microarray. The results suggest that CRP activates the expression of former four sRNAs but represses the latter two sRNAs.

**Supplemental Table S1.** Targets of CRP-regulated sRNA also have CRP binding sites.

| Group | Number of sRNA | CRP binding sites | Differential expression | Target with CRP binding sites | P value (hypergeometric test) based on RegulonDB | P value (hypergeometric test) based on HMM prediction |
| --- | --- | --- | --- | --- | --- | --- |
| Total sRNA | 80 | (37/79) | (36/79) | 71% (61/86) | $P < 3 \times 10^{-68}$ | $P < 2 \times 10^{-20}$ |
| Direct | 25 | Presence | Yes | 84% (40/48) | $P < 3 \times 10^{-49}$ | $P < 4 \times 10^{-18}$ |
| Conditional | 12 | Presence | No | 56% (9/16) | $P < 3 \times 10^{-9}$ | $P < 4 \times 10^{-3}$ |
| Indirect | 16 | Absence | Yes | 86% (6/7) | $P < 3 \times 10^{-8}$ | $P < 1 \times 10^{-3}$ |
| Independent | 27 | Absence | No | 44% (7/16) | $P < 2 \times 10^{-6}$ | $P < 4 \times 10^{-2}$ |

**Supplemental Table S3.**
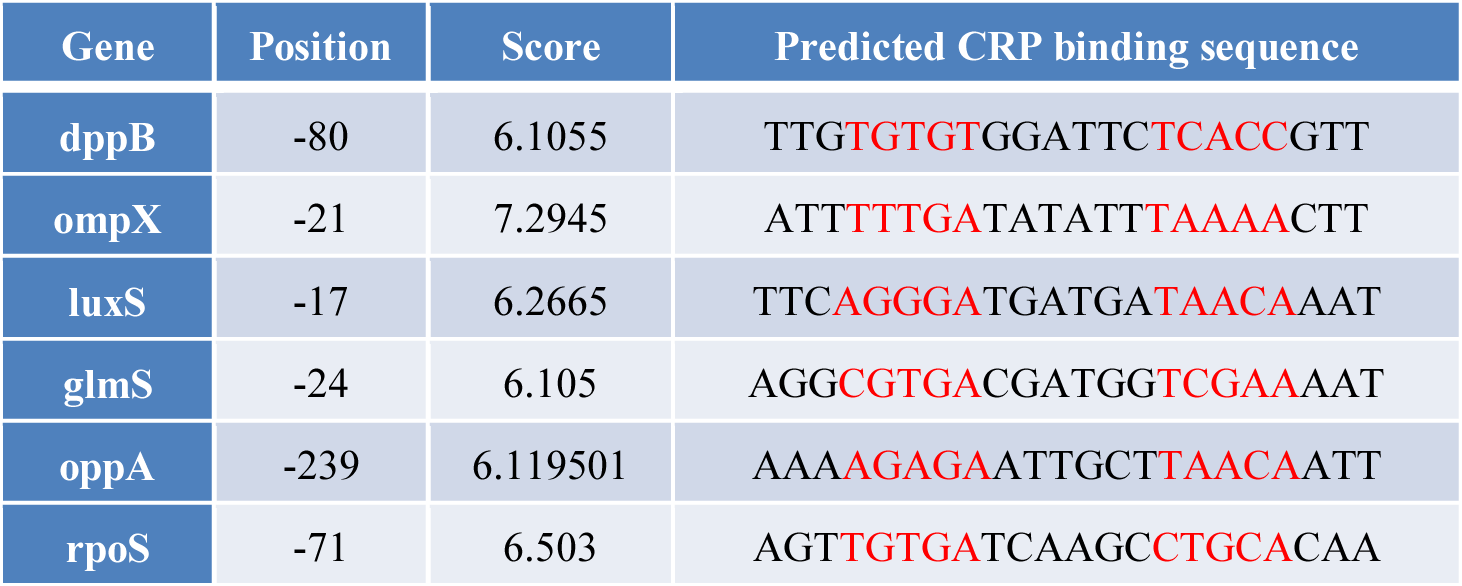
The predicted CRP binding sites in the promoter regions of some sRNA target genes. The table listed the location, score, as well as the sequence of predicted CRP binding sites. The CRP binding ability was further confirmed by EMSA. Synthesized oligonucleotides were incubated with cAMP (200μM) and CRP at various concentrations (from 0 to 200 nM) in the buffer.

## REFERENCES

1. Milo, R., Shen-Orr, S., Itzkovitz, S., Kashtan, N., Chklovskii, D. and Alon, U. (2002) Network motifs: simple building blocks of complex networks. Science, 298, 824–827.

2. Linding, R., Jensen, L.J., Ostheimer, G.J., van Vugt, M.A., Jorgensen, C., Miron, I.M., Diella, F., Colwill, K., Taylor, L., Elder, K., et al. (2007) Systematic discovery of in vivo phosphorylation networks. Cell, 129, 1415–1426.

3. Mason, S.D. and Joyce, J.A. (2011) Proteolytic networks in cancer. Trends in cell biology, 21, 228–237.

4. Ray, J.C., Tabor, J.J. and Igoshin, O.A. (2011) Non-transcriptional regulatory processes shape transcriptional network dynamics. Nature reviews. Microbiology, 9, 817–828.

5. Waters, L.S. and Storz, G. (2009) Regulatory RNAs in bacteria. Cell, 136, 615–628.

6. Majdalani, N., Vanderpool, C.K. and Gottesman, S. (2005) Bacterial small RNA regulators. Crit Rev Biochem Mol Biol, 40, 93–113.

7. Repoila, F. and Darfeuille, F. (2009) Small regulatory non-coding RNAs in bacteria: physiology and mechanistic aspects. Biology of the cell / under the auspices of the European Cell Biology Organization, 101, 117–131.

8. Suzuki, K., Wang, X., Weilbacher, T., Pernestig, A.K., Melefors, O., Georgellis, D., Babitzke, P. and Romeo, T. (2002) Regulatory circuitry of the CsrA/CsrB and BarA/UvrY systems of Escherichia coli. J Bacteriol, 184, 5130–5140.

9. Jonas, K. and Melefors, O. (2009) The Escherichia coli CsrB and CsrC small RNAs are strongly induced during growth in nutrient-poor medium. FEMS Microbiol Lett, 297, 80–86.

10. Reichenbach, B., Maes, A., Kalamorz, F., Hajnsdorf, E. and Gorke, B. (2008) The small RNA GlmY acts upstream of the sRNA GlmZ in the activation of glmS expression and is subject to regulation by polyadenylation in Escherichia coli. Nucleic acids research, 36, 2570–2580.

11. van Hijum, S.A., Medema, M.H. and Kuipers, O.P. (2009) Mechanisms and evolution of control logic in prokaryotic transcriptional regulation. Microbiology and molecular biology reviews : MMBR, 73, 481–509, Table of Contents.

12. Thomason, M.K., Fontaine, F., De Lay, N. and Storz, G. (2012) A small RNA that regulates motility and biofilm formation in response to changes in nutrient availability in Escherichia coli. Molecular microbiology, 84, 17–35.

13. Pulvermacher, S.C., Stauffer, L.T. and Stauffer, G.V. (2009) Role of the sRNA GcvB in regulation of cycA in Escherichia coli. Microbiology, 155, 106–114.

14. Weilbacher, T., Suzuki, K., Dubey, A.K., Wang, X., Gudapaty, S., Morozov, I., Baker, C.S., Georgellis, D., Babitzke, P. and Romeo, T. (2003) A novel sRNA component of the carbon storage regulatory system of Escherichia coli. Mol Microbiol, 48, 657–670.

15. Hussein, R. and Lim, H.N. (2012) Direct comparison of small RNA and transcription factor signaling. Nucleic acids research.

16. Mehta, P., Goyal, S. and Wingreen, N.S. (2008) A quantitative comparison of sRNA-based and protein-based gene regulation. Molecular systems biology, 4, 221.

17. Masse, E., Escorcia, F.E. and Gottesman, S. (2003) Coupled degradation of a small regulatory RNA and its mRNA targets in Escherichia coli. Gene Dev, 17, 2374–2383.

18. Eddy, S.R. (2001) Non-coding RNA genes and the modern RNA world. Nature reviews. Genetics, 2, 919–929.

19. Li, Z., Gong, X., Joshi, V.H. and Li, M. (2005) Co-evolution of tRNA 3’ trailer sequences with 3’ processing enzymes in bacteria. RNA, 11, 567–577.

20. Zwieb, C., van Nues, R.W., Rosenblad, M.A., Brown, J.D. and Samuelsson, T. (2005) A nomenclature for all signal recognition particle RNAs. RNA, 11, 7–13.

21. Liu, D., Chang, X., Liu, Z., Chen, L. and Wang, R. (2011) Bistability and oscillations in gene regulation mediated by small noncoding RNAs. PloS one, 6, e17029.

22. Tu, K.C., Waters, C.M., Svenningsen, S.L. and Bassler, B.L. (2008) A small-RNA-mediated negative feedback loop controls quorum-sensing dynamics in Vibrio harveyi. Molecular microbiology, 70, 896–907.

23. Shimoni, Y., Friedlander, G., Hetzroni, G., Niv, G., Altuvia, S., Biham, O. and Margalit, H. (2007) Regulation of gene expression by small non-coding RNAs: a quantitative view. Molecular systems biology, 3, 138.

24. Busby, S. and Ebright, R.H. (1999) Transcription activation by catabolite activator protein (CAP). Journal of molecular biology, 293, 199–213.

25. Hudson, J.M. and Fried, M.G. (1990) Co-operative interactions between the catabolite gene activator protein and the lac repressor at the lactose promoter. Journal of molecular biology, 214, 381–396.

26. Kiley, P.J. and Beinert, H. (1998) Oxygen sensing by the global regulator, FNR: the role of the iron-sulfur cluster. FEMS microbiology reviews, 22, 341–352.

27. Mills, S.A. and Marletta, M.A. (2005) Metal binding characteristics and role of iron oxidation in the ferric uptake regulator from Escherichia coli. Biochemistry, 44, 13553–13559.

28. Forst, S., Delgado, J. and Inouye, M. (1989) Phosphorylation of OmpR by the osmosensor EnvZ modulates expression of the ompF and ompC genes in Escherichia coli. Proc Natl Acad Sci U S A, 86, 6052–6056.

29. Thomas, J.G. and Baneyx, F. (1998) Roles of the Escherichia coli small heat shock proteins IbpA and IbpB in thermal stress management: comparison with ClpA, ClpB, and HtpG In vivo. Journal of bacteriology, 180, 5165–5172.

30. Tetsch, L., Koller, C., Donhofer, A. and Jung, K. (2011) Detection and function of an intramolecular disulfide bond in the pH-responsive CadC of Escherichia coli. BMC microbiology, 11, 74.

31. Tsang, J., Zhu, J. and van Oudenaarden, A. (2007) MicroRNA-mediated feedback and feedforward loops are recurrent network motifs in mammals. Molecular cell, 26, 753–767.

32. Neph, S., Stergachis, A.B., Reynolds, A., Sandstrom, R., Borenstein, E. and Stamatoyannopoulos, J.A. (2012) Circuitry and dynamics of human transcription factor regulatory networks. Cell, 150, 1274–1286.

33. Alon, U. (2007) Network motifs: theory and experimental approaches. Nature reviews. Genetics, 8, 450–461.

34. Rodrigo, G. and Elena, S.F. (2011) Structural discrimination of robustness in transcriptional feedforward loops for pattern formation. PloS one, 6, e16904.

35. Osella, M., Bosia, C., Cora, D. and Caselle, M. (2011) The role of incoherent microRNA-mediated feedforward loops in noise buffering. PLoS computational biology, 7, e1001101.

36. Goentoro, L., Shoval, O., Kirschner, M.W. and Alon, U. (2009) The incoherent feedforward loop can provide fold-change detection in gene regulation. Molecular cell, 36, 894–899.

37. Mangan, S. and Alon, U. (2003) Structure and function of the feed-forward loop network motif. Proceedings of the National Academy of Sciences of the United States of America, 100, 11980–11985.

38. Beisel, C.L. and Storz, G. (2011) The base-pairing RNA spot 42 participates in a multioutput feedforward loop to help enact catabolite repression in Escherichia coli. Molecular cell, 41, 286–297.

39. Nolan, T., Hands, R.E. and Bustin, S.A. (2006) Quantification of mRNA using real-time RT-PCR. Nature protocols, 1, 1559–1582.

40. Keseler, I.M., Collado-Vides, J., Santos-Zavaleta, A., Peralta-Gil, M., Gama-Castro, S., Muniz-Rascado, L., Bonavides-Martinez, C., Paley, S., Krummenacker, M., Altman, T. et al. (2011) EcoCyc: a comprehensive database of Escherichia coli biology. Nucleic acids research, 39, D583–590.

41. Chen, Y.G., Lin, Y.C.D., Luo, Y.J., Cai, X.X., Qiu, P., Cui, S.D., Wang, Z., Huang, H.Y. and Huang, H.D. (2023) Quantitative model for genome-wide cyclic AMP receptor protein binding site identification and characteristic analysis. Brief Bioinform, 24.

42. Chen, Y.G., Mao, R.B., Xu, J.T., Huang, Y.X., Xu, J.Y., Cui, S.D., Zhu, Z.H., Ji, X., Huang, S.H., Huang, Y.Z. et al. (2024) A Causal Regulation Modeling Algorithm for Temporal Events with Application to ’s Aerobic to Anaerobic Transition. Int J Mol Sci, 25.

43. Yang, C.D., Huang, H.Y., Shrestha, S., Chen, Y.H., Huang, H.D. and Tseng, C.P. (2018) Large-Scale Functional Analysis of CRP-Mediated Feed-Forward Loops. Int J Mol Sci, 19.

44. Huang, H.Y., Chang, H.Y., Chou, C.H., Tseng, C.P., Ho, S.Y., Yang, C.D., Ju, Y.W. and Huang, H.D. (2009) sRNAMap: genomic maps for small non-coding RNAs, their regulators and their targets in microbial genomes. Nucleic acids research, 37, D150–154.

45. Gama-Castro, S., Salgado, H., Peralta-Gil, M., Santos-Zavaleta, A., Muniz-Rascado, L., Solano-Lira, H., Jimenez-Jacinto, V., Weiss, V., Garcia-Sotelo, J.S., Lopez-Fuentes, A. et al. (2011) RegulonDB version 7.0: transcriptional regulation of Escherichia coli K-12 integrated within genetic sensory response units (Gensor Units). Nucleic acids research, 39, D98–105.

46. Wu, T., Wang, J., Liu, C., Zhang, Y., Shi, B., Zhu, X., Zhang, Z., Skogerbo, G., Chen, L., Lu, H. et al. (2006) NPInter: the noncoding RNAs and protein related biomacromolecules interaction database. Nucleic Acids Res, 34, D150–152.

47. Cao, Y., Wu, J., Liu, Q., Zhao, Y., Ying, X., Cha, L., Wang, L. and Li, W. (2010) sRNATarBase: a comprehensive database of bacterial sRNA targets verified by experiments. RNA, 16, 2051–2057.

48. Mandin, P. and Gottesman, S. (2010) Integrating anaerobic/aerobic sensing and the general stress response through the ArcZ small RNA. The EMBO journal, 29, 3094–3107.

49. Durand, S. and Storz, G. (2010) Reprogramming of anaerobic metabolism by the FnrS small RNA. Mol Microbiol, 75, 1215–1231.

50. Vanderpool, C.K. (2011) Combined experimental and computational strategies define an expansive regulon for GcvB small RNA. Mol Microbiol, 81, 1129–1132.

51. Antal, M., Bordeau, V., Douchin, V. and Felden, B. (2005) A small bacterial RNA regulates a putative ABC transporter. J Biol Chem, 280, 7901–7908.

52. Papenfort, K., Pfeiffer, V., Mika, F., Lucchini, S., Hinton, J.C. and Vogel, J. (2006) SigmaE-dependent small RNAs of Salmonella respond to membrane stress by accelerating global omp mRNA decay. Mol Microbiol, 62, 1674–1688.

53. Johansen, J., Eriksen, M., Kallipolitis, B. and Valentin-Hansen, P. (2008) Down-regulation of outer membrane proteins by noncoding RNAs: unraveling the cAMP-CRP- and sigmaE-dependent CyaR-ompX regulatory case. Journal of molecular biology, 383, 1–9.

54. Hobbs, E.C., Astarita, J.L. and Storz, G. (2010) Small RNAs and small proteins involved in resistance to cell envelope stress and acid shock in Escherichia coli: analysis of a bar-coded mutant collection. J Bacteriol, 192, 59–67.

55. Liu, M.Y., Gui, G., Wei, B., Preston, J.F., 3rd, Oakford, L., Yuksel, U., Giedroc, D.P. and Romeo, T. (1997) The RNA molecule CsrB binds to the global regulatory protein CsrA and antagonizes its activity in Escherichia coli. J Biol Chem, 272, 17502–17510.

56. Zheng, D., Constantinidou, C., Hobman, J.L. and Minchin, S.D. (2004) Identification of the CRP regulon using in vitro and in vivo transcriptional profiling. Nucleic acids research, 32, 5874–5893.

57. Gosset, G., Zhang, Z., Nayyar, S., Cuevas, W.A. and Saier, M.H., Jr. (2004) Transcriptome analysis of Crp-dependent catabolite control of gene expression in Escherichia coli. Journal of bacteriology, 186, 3516–3524.

58. Valentin-Hansen, P., Holst, B., Sogaard-Andersen, L., Martinussen, J., Nesvera, J. and Douthwaite, S.R. (1991) Design of cAMP-CRP-activated promoters in Escherichia coli. Molecular microbiology, 5, 433–437.

59. Smirnov Iu, V. and Lisenkov, A.F. (1986) [Construction of the hybrid crp-lac operon and study of the role of CRP-cAMP complex in its regulation in Escherichia coli]. Genetika, 22, 576–583.

60. Cossart, P. and Gicquel-Sanzey, B. (1985) Regulation of expression of the crp gene of Escherichia coli K-12: in vivo study. Journal of bacteriology, 161, 454–457.

61. Hantke, K., Winkler, K. and Schultz, J.E. (2011) Escherichia coli exports cyclic AMP via TolC. Journal of bacteriology, 193, 1086–1089.

62. Dubey, A.K., Baker, C.S., Romeo, T. and Babitzke, P. (2005) RNA sequence and secondary structure participate in high-affinity CsrA-RNA interaction. RNA, 11, 1579–1587.

63. Baker, C.S., Morozov, I., Suzuki, K., Romeo, T. and Babitzke, P. (2002) CsrA regulates glycogen biosynthesis by preventing translation of glgC in Escherichia coli. Mol Microbiol, 44, 1599–1610.

64. Baker, C.S., Eory, L.A., Yakhnin, H., Mercante, J., Romeo, T. and Babitzke, P. (2007) CsrA inhibits translation initiation of Escherichia coli hfq by binding to a single site overlapping the Shine-Dalgarno sequence. J Bacteriol, 189, 5472–5481.

65. Yang, H., Liu, M.Y. and Romeo, T. (1996) Coordinate genetic regulation of glycogen catabolism and biosynthesis in Escherichia coli via the CsrA gene product. J Bacteriol, 178, 1012–1017.

66. Hengge-Aronis, R. and Fischer, D. (1992) Identification and molecular analysis of glgS, a novel growth-phase-regulated and rpoS-dependent gene involved in glycogen synthesis in Escherichia coli. Mol Microbiol, 6, 1877–1886.

67. Masse, E., Majdalani, N. and Gottesman, S. (2003) Regulatory roles for small RNAs in bacteria. Current opinion in microbiology, 6, 120–124.

68. Hwang, W., Arluison, V. and Hohng, S. (2011) Dynamic competition of DsrA and rpoS fragments for the proximal binding site of Hfq as a means for efficient annealing. Nucleic acids research, 39, 5131–5139.

69. Williams, R.M. and Rimsky, S. (1997) Molecular aspects of the E. coli nucleoid protein, H-NS: a central controller of gene regulatory networks. FEMS microbiology letters, 156, 175–185.

70. Schauder, S., Shokat, K., Surette, M.G. and Bassler, B.L. (2001) The LuxS family of bacterial autoinducers: biosynthesis of a novel quorum-sensing signal molecule. Molecular microbiology, 41, 463–476.

71. Papenfort, K., Pfeiffer, V., Lucchini, S., Sonawane, A., Hinton, J.C. and Vogel, J. (2008) Systematic deletion of Salmonella small RNA genes identifies CyaR, a conserved CRP-dependent riboregulator of OmpX synthesis. Molecular microbiology, 68, 890–906.

72. Gunasekera, A., Ebright, Y.W. and Ebright, R.H. (1992) DNA sequence determinants for binding of the Escherichia coli catabolite gene activator protein. J Biol Chem, 267, 14713–14720.

73. Hollands, K., Busby, S.J. and Lloyd, G.S. (2007) New targets for the cyclic AMP receptor protein in the Escherichia coli K-12 genome. FEMS Microbiol Lett, 274, 89–94.

74. Brikun, I., Suziedelis, K., Stemmann, O., Zhong, R., Alikhanian, L., Linkova, E., Mironov, A. and Berg, D.E. (1996) Analysis of CRP-CytR interactions at the Escherichia coli udp promoter. Journal of bacteriology, 178, 1614–1622.

75. Hermsen, R., Tans, S. and ten Wolde, P.R. (2006) Transcriptional regulation by competing transcription factor modules. PLoS Comput Biol, 2, e164.

76. Beisel, C.L. and Storz, G. (2010) Base pairing small RNAs and their roles in global regulatory networks. FEMS microbiology reviews, 34, 866–882.

77. Mangan, S., Itzkovitz, S., Zaslaver, A. and Alon, U. (2006) The incoherent feed-forward loop accelerates the response-time of the gal system of Escherichia coli. Journal of molecular biology, 356, 1073–1081.

78. Kline, E.L., Brown, C.S., Bankaitis, V., Montefiori, D.C. and Craig, K. (1980) Metabolite gene regulation of the L-arabinose operon in Escherichia coli with indoleacetic acid and other indole derivatives. Proc Natl Acad Sci U S A, 77, 1768–1772.

79. Kline, E.L., Bankaitis, V.A., Brown, C.S. and Montefiori, D.C. (1980) Metabolite gene regulation: imidazole and imidazole derivatives which circumvent cyclic adenosine 3’,5’-monophosphate in induction of the Escherichia coli L-arabinose operon. J Bacteriol, 141, 770–778.

80. Sledjeski, D.D., Gupta, A. and Gottesman, S. (1996) The small RNA, DsrA, is essential for the low temperature expression of RpoS during exponential growth in Escherichia coli. The EMBO journal, 15, 3993–4000.

81. Wassarman, K.M., Repoila, F., Rosenow, C., Storz, G. and Gottesman, S. (2001) Identification of novel small RNAs using comparative genomics and microarrays. Genes & development, 15, 1637–1651.

82. Lange, R. and Hengge-Aronis, R. (1994) The cellular concentration of the sigma S subunit of RNA polymerase in Escherichia coli is controlled at the levels of transcription, translation, and protein stability. Genes & development, 8, 1600–1612.

83. Majdalani, N., Cunning, C., Sledjeski, D., Elliott, T. and Gottesman, S. (1998) DsrA RNA regulates translation of RpoS message by an anti-antisense mechanism, independent of its action as an antisilencer of transcription. Proceedings of the National Academy of Sciences of the United States of America, 95, 12462–12467.

84. Lease, R.A., Cusick, M.E. and Belfort, M. (1998) Riboregulation in Escherichia coli: DsrA RNA acts by RNA:RNA interactions at multiple loci. Proceedings of the National Academy of Sciences of the United States of America, 95, 12456–12461.

85. Repoila, F. and Gottesman, S. (2001) Signal transduction cascade for regulation of RpoS: temperature regulation of DsrA. Journal of bacteriology, 183, 4012–4023.

86. Kalamorz, F., Reichenbach, B., Marz, W., Rak, B. and Gorke, B. (2007) Feedback control of glucosamine-6-phosphate synthase GlmS expression depends on the small RNA GlmZ and involves the novel protein YhbJ in Escherichia coli. Molecular microbiology, 65, 1518–1533.

87. Chang, D.E., Smalley, D.J., Tucker, D.L., Leatham, M.P., Norris, W.E., Stevenson, S.J., Anderson, A.B., Grissom, J.E., Laux, D.C., Cohen, P.S. et al. (2004) Carbon nutrition of Escherichia coli in the mouse intestine. Proceedings of the National Academy of Sciences of the United States of America, 101, 7427–7432.

88. Soper, T., Mandin, P., Majdalani, N., Gottesman, S. and Woodson, S.A. (2010) Positive regulation by small RNAs and the role of Hfq. Proceedings of the National Academy of Sciences of the United States of America, 107, 9602–9607.

89. Moller, T., Franch, T., Hojrup, P., Keene, D.R., Bachinger, H.P., Brennan, R.G. and Valentin-Hansen, P. (2002) Hfq: a bacterial Sm-like protein that mediates RNA-RNA interaction. Molecular cell, 9, 23–30.

90. Romeo, T. (1998) Global regulation by the small RNA-binding protein CsrA and the non-coding RNA molecule CsrB. Mol Microbiol, 29, 1321–1330.

## REFERENCE

Baba, T., Ara, T., Hasegawa, M., Takai, Y., Okumura, Y., Baba, M., Datsenko, K.A., Tomita, M., Wanner, B.L., and Mori, H. (2006). Construction of Escherichia coli K-12 in-frame, single-gene knockout mutants: the Keio collection. Mol Syst Biol 2, 2006 0008.

Becskei, A., Seraphin, B., and Serrano, L. (2001). Positive feedback in eukaryotic gene networks: cell differentiation by graded to binary response conversion. Embo J 20, 2528–2535.

Elowitz, M.B., and Leibler, S. (2000). A synthetic oscillatory network of transcriptional regulators. Nature 403, 335–338.

Ferrell, J.E., and Machleder, E.M. (1998). The biochemical basis of an all-or-none cell fate switch in Xenopus oocytes. Science 280, 895–898.

Finn, R.D., Clements, J., and Eddy, S.R. (2011). HMMER web server: interactive sequence similarity searching. Nucleic Acids Res 39 *Suppl 2*, W29–37.

Fukami-Kobayashi, K., and Saito, N. (2002). [How to make good use of CLUSTALW]. Tanpakushitsu Kakusan Koso 47, 1237–1239.

Gardner, T.S., Cantor, C.R., and Collins, J.J. (2000). Construction of a genetic toggle switch in Escherichia coli. Nature 403, 339–342.

Gunasekera, A., Ebright, Y.W., and Ebright, R.H. (1992). DNA sequence determinants for binding of the Escherichia coli catabolite gene activator protein. J Biol Chem 267, 14713–14720.

Hantke, K., Winkler, K., and Schultz, J.E. (2011). Escherichia coli exports cyclic AMP via TolC. Journal of bacteriology 193, 1086–1089.

Kel, A.E., Gossling, E., Reuter, I., Cheremushkin, E., Kel-Margoulis, O.V., and Wingender, E. (2003). MATCH: A tool for searching transcription factor binding sites in DNA sequences. Nucleic Acids Res 31, 3576–3579.

Keseler, I.M., Collado-Vides, J., Santos-Zavaleta, A., Peralta-Gil, M., Gama-Castro, S., Muniz-Rascado, L., Bonavides-Martinez, C., Paley, S., Krummenacker, M., Altman, T., et al. (2011). EcoCyc: a comprehensive database of Escherichia coli biology. Nucleic Acids Res 39, D583–590.

Larkin, M.A., Blackshields, G., Brown, N.P., Chenna, R., McGettigan, P.A., McWilliam, H., Valentin, F., Wallace, I.M., Wilm, A., Lopez, R., et al. (2007). Clustal W and Clustal X version 2.0. Bioinformatics 23, 2947–2948.

Legewie, S., Dienst, D., Wilde, A., Herzel, H., and Axmann, I.M. (2008). Small RNAs establish delays and temporal thresholds in gene expression. Biophys J 95, 3232–3238.

Levine, E., Zhang, Z., Kuhlman, T., and Hwa, T. (2007). Quantitative characteristics of gene regulation by small RNA. Plos Biol 5, 1998–2010.

Lin, H.H., Hsu, C.C., Yang, C.D., Ju, Y.W., Chen, Y.P., and Tseng, C.P. (2011). Negative effect of glucose on ompA mRNA stability: a potential role of cyclic AMP in the repression of hfq in Escherichia coli. Journal of bacteriology 193, 5833–5840.

Rosenfeld, N., Elowitz, M.B., and Alon, U. (2002). Negative autoregulation speeds the response times of transcription networks. J Mol Biol 323, 785–793.

Shimoni, Y., Friedlander, G., Hetzroni, G., Niv, G., Altuvia, S., Biham, O., and Margalit, H. (2007). Regulation of gene expression by small non-coding RNAs: a quantitative view. Molecular systems biology 3, 138.

Soper, T., Mandin, P., Majdalani, N., Gottesman, S., and Woodson, S.A. (2010). Positive regulation by small RNAs and the role of Hfq. P Natl Acad Sci USA 107, 9602–9607.

## Reference

Bleris, L., Xie, Z., Glass, D., Adadey, A., Sontag, E., and Benenson, Y. (2011). Synthetic incoherent feedforward circuits show adaptation to the amount of their genetic template. Molecular systems biology 7.

Cao, Y., Wu, J., Liu, Q., Zhao, Y., Ying, X., Cha, L., Wang, L., and Li, W. (2010). sRNATarBase: a comprehensive database of bacterial sRNA targets verified by experiments. RNA 16, 2051–2057.

Crooks, G.E., Hon, G., Chandonia, J.M., and Brenner, S.E. (2004). WebLogo: a sequence logo generator. Genome research 14, 1188–1190.

Ellis, T., Wang, X., and Collins, J.J. (2009). Diversity-based, model-guided construction of synthetic gene networks with predicted functions. Nat Biotechnol 27, 465–471.

Gama-Castro, S., Salgado, H., Peralta-Gil, M., Santos-Zavaleta, A., Muniz-Rascado, L., Solano-Lira, H., Jimenez-Jacinto, V., Weiss, V., Garcia-Sotelo, J.S., Lopez-Fuentes, A., et al. (2011). RegulonDB version 7.0: transcriptional regulation of Escherichia coli K-12 integrated within genetic sensory response units (Gensor Units). Nucleic acids research 39, D98–105.

Goentoro, L., Shoval, O., Kirschner, M.W., and Alon, U. (2009). The incoherent feedforward loop can provide fold-change detection in gene regulation. Molecular cell 36, 894–899.

Kaplan, S., Bren, A., Dekel, E., and Alon, U. (2008). The incoherent feed-forward loop can generate non-monotonic input functions for genes. Molecular systems biology 4.

Kuttykrishnan, S., Sabina, J., Langton, L.L., Johnston, M., and Brent, M.R. (2010). A quantitative model of glucose signaling in yeast reveals an incoherent feed forward loop leading to a specific, transient pulse of transcription. P Natl Acad Sci USA 107, 16743–16748.

Legewie, S., Dienst, D., Wilde, A., Herzel, H., and Axmann, I.M. (2008a). Small RNAs establish delays and temporal thresholds in gene expression. Biophysical journal 95, 3232–3238.

Legewie, S., Dienst, D., Wilde, A., Herzel, H., and Axmann, I.M. (2008b). Small RNAs establish delays and temporal thresholds in gene expression. Biophys J 95, 3232–3238.

Macia, J., Widder, S., and Sole, R. (2009). Specialized or flexible feed-forward loop motifs: a question of topology. BMC systems biology 3, 84.

Osella, M., Bosia, C., Cora, D., and Caselle, M. (2011). The role of incoherent microRNA-mediated feedforward loops in noise buffering. Plos Comput Biol 7, e1001101.

Rodrigo, G., and Elena, S.F. (2011). Structural discrimination of robustness in transcriptional feedforward loops for pattern formation. PloS one 6, e16904.

Sledjeski, D.D., Gupta, A., and Gottesman, S. (1996). The small RNA, DsrA, is essential for the low temperature expression of RpoS during exponential growth in Escherichia coli. The EMBO journal 15, 3993–4000.

Wu, T., Wang, J., Liu, C., Zhang, Y., Shi, B., Zhu, X., Zhang, Z., Skogerbo, G., Chen, L., Lu, H., et al. (2006). NPInter: the noncoding RNAs and protein related biomacromolecules interaction database. Nucleic Acids Res 34, D150–152.

